# Lupus auto-antibodies act as positive allosteric modulators at NMDA receptors and induce spatial memory deficits

**DOI:** 10.1101/791715

**Authors:** Kelvin Chan, Jacquelyn Nestor, Tomás S. Huerta, Noele Certain, Gabrielle Moody, Czeslawa Kowal, Patricio T. Huerta, Bruce T. Volpe, Betty Diamond, Lonnie P. Wollmuth

**Author notes:** These authors contributed equally to this manuscript (Kelvin Chan, Jacquelyn Nestor). Co-senior and corresponding authors; Address for correspondence: Dr. Lonnie P. Wollmuth, Depts. of Neurobiology & Behavior and Biochemistry & Cell Biology, Center for Nervous System Disorders, Stony Brook University, Stony Brook, New York 11794-5230, Dr. Betty Diamond, Center for Autoimmune, Musculoskeletal and Hematopoietic Diseases, Feinstein Institute for Medical Research, Northwell Health, Manhasset, New York 11030.

## Abstract

Patients with Systemic lupus erythematosus (SLE) experience various peripheral and central nervous system manifestations including spatial memory impairment. A subset of auto-antibodies (DNRAbs) cross-react with the GluN2A and GluN2B subunits of the NMDA receptor (NMDAR). We find that these DNRAbs act as positive allosteric modulators on NMDARs with GluN2A-containing NMDARs, even those containing a single GluN2A subunit, exhibiting a much greater sensitivity to DNRAbs than those with exclusively GluN2B. Accordingly, GluN2A-specific antagonists provide greater protection from DNRAb-mediated neuronal cell death than GluN2B antagonists. Using transgenic mice to perturb expression of either GluN2A or GluN2B *in vivo*, we find that DNRAb-mediated disruption of spatial memory characterized by early neuronal cell death and subsequent microglia-dependent pathologies requires GluN2A-containing NMDARs. Our results indicate that GluN2A-specific antagonists or negative allosteric modulators are strong candidates to treat SLE patients with nervous system dysfunction.

## INTRODUCTION

Systemic lupus erythematosus (SLE) is an autoimmune disease characterized by the presence of autoantibodies directed against multiple self-antigens, including DNA^1^. These autoantibodies affect multiple organ systems such that SLE patients experience arthritis, renal disease, anemia, rashes, and neuropsychiatric symptoms including memory disorders and spatial memory impairment^2, 3, 4^. The prevalence of diffuse nervous system disorders is reported from 20-90% depending on the particular functional assessment^5, 6, 7^. These cognitive defects in both clinical and pre-clinical conditions are often associated with DNRAb, anti-double stranded DNA (dsDNA) antibodies with cross-reactivity to NMDA receptors (NMDAR)^8, 9, 10, 11, 12, 13, 14^.

The role of DNRAbs in contributing to neuropsychiatric symptoms in SLE have largely been studied in mice models that endogenously synthesize DNRAbs and in mice exposed to patient-derived DNRAbs^8, 10, 11^. Patient-derived DNRAbs are IgG1 antibodies cloned from patient B cells that display reactivity to dsDNA and NMDARs^15, 16^. Specific regions of the brain in mice are targeted based on the experimental method used to trigger blood brain barrier permeability – lipopolysaccharide (LPS) causes DNRAbs to deposit in the hippocampus while epinephrine causes DNRAbs to deposit in the amygdala^11, 17^. Non-invasive imaging of SLE patients have revealed hippocampal atrophy and parahippocampal microstructural defects, conferring an advantage to using LPS in disease models^4, 18^.

LPS-treated mice immunogenized to generate DNRAbs and patient-derived DNRAbs display a gamut of pathologies in the hippocampus: aberrant excitatory signaling, apoptosis, dendritic pruning, and microglial activation^10, 11, 19^. These mice also display expanded place fields in the hippocampus and defects in spatial memory^2, 19^. These studies are essential to defining the pathology of the neuropsychiatric component of SLE, but do not define the specific NMDARs that mediate these effects. This information is critical to potentially develop therapies to treat and prevent neuropsychiatric symptoms associated with DNRAbs.

NMDARs are ionotropic glutamate receptors that are central to excitatory synaptic transmission in the brain. NMDARs are heterotetramers composed of two obligate GluN1 subunits and typically two GluN2 subunits of the same or different subtype (GluN2A, B, C or D)^20, 21^. DNRAbs bind to both GluN2A and GluN2B subunits, with the epitope including a pentapeptide consensus sequence, DWEYS^10, 11^. This antigenic target for DNRAbs is in the extracellularly located amino-terminal domain (ATD) of GluN2, an allosteric hub for modulating NMDAR function^21, 22^. NMDARs containing GluN2A or GluN2B have distinct physiological, pharmacological, and signaling properties^22^. Nevertheless, the contribution of the different GluN2 subunits to the SLE-associated neuropathologies is unclear. Knockout studies in mice have suggested GluN2A to be the primary target for DNRAb-mediated adult and fetal neuronal cell death^23^, though the evidence was limited. Specific inhibitors of GluN2B also reduced DNRAb-mediated cell death, suggesting a significant contribution of GluN2B to the DNRAb-mediated phenotype^10^. All these results are ambiguous because of uncertainty of concentrations and technical limitations.

Here, we use a combination of heterologous expression systems and animal models to show subunit-specific susceptibility to DNRAb-mediated pathological effects. For heterologous expression, we tested a DNRAb (G11 and its B1 isotype control) derived from a human patient (see Materials and Methods) in concentrations relevant to patient CSF levels^10^. We find that DNRAbs act as positive allosteric modulators (PAMs) at both GluN2A- and GluN2B-containing NMDARs, but that GluN2A-containing receptors have a much higher intrinsic sensitivity to DNRAb-mediated potentiation than GluN2B-containing NMDARs. Using a heterologous expression system to express tri-heteromeric GluN1/GluN2A/GluN2B, we find that a single GluN2A subunit confers high sensitivity to DNRAbs. We find that mice with a forebrain deletion of the GluN2B subunit display the full spectrum of DNRAb-mediated pathology, including acute loss of hippocampal CA1 neurons, dendritic abnormalities in surviving neurons, microglia activation, defective place cell fields, and impaired spatial memory. Conversely, GluN2A knockout mice are protected from the effects of DNRAbs. Thus, our work identifies the GluN2A subunit as the central mediator of NMDAR-associated nervous system pathology in SLE, supporting the use of GluN2A-specific negative allosteric modulators to treat SLE patients with brain dysfunction.

## RESULTS

### GluN2A-containing NMDARs are significantly more sensitive to DNRAbs than GluN2B-containing receptors

DNRAbs bind to both GluN2A and GluN2B subunits^2, 10^. To begin to address how these subunits contribute to DNRAb-induced phenotypes, we characterized the effect of a SLE patient-derived monoclonal DNRAb, G11, on heterologously expressed NMDARs composed of human NMDAR subunits, either hGluN1a/hGluN2A (hN1/hN2A) or hGluN1a/hGluN2B (hN1/hN2B). In parallel with antibody titers found in SLE patient CSF samples^10^, we test the DNRAbs at concentrations from 1-100 μg/ml. For macroscopic or whole cell currents, exposure of hN1/hN2A NMDARs to 10 μg/ml of G11 (green traces) strongly potentiated glutamate-gated current amplitudes compared to baseline (Figure 1A). In contrast, a similar exposure to N2B-containing receptors had no effect on current amplitudes (Figure 1B). A matched concentration of an isotype control antibody, B1 (gray traces), had no effect on current amplitudes for either N2A- or N2B-containing receptors.

**Figure 1.**
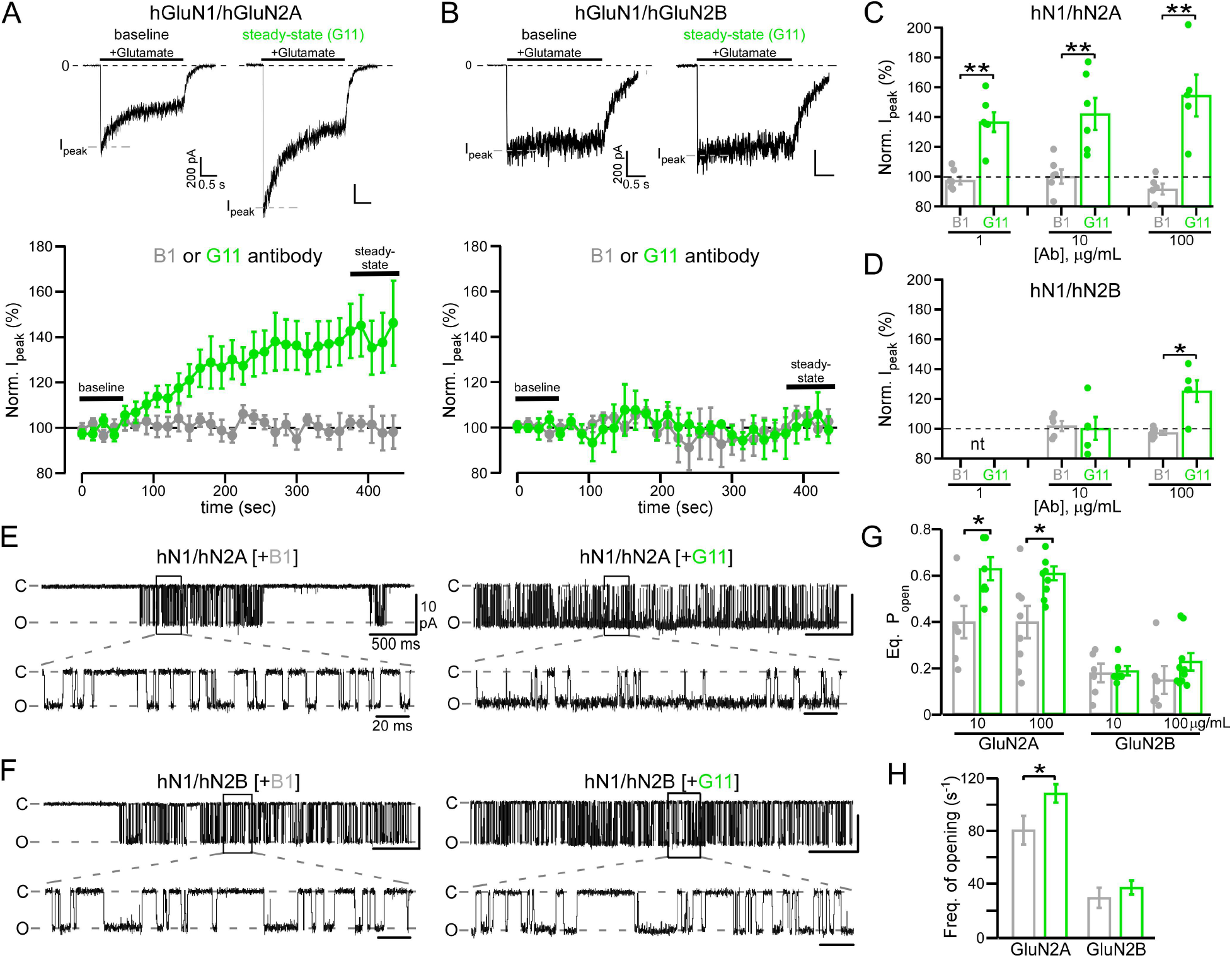
Differential sensitivity of N2A- and N2B-containing NMDARs to DNRAbs. (**A & B**) Moderate pathophysiological levels of DNRAbs (10 μg/mL) potentiate glutamate-activated currents in N2A-containing, but not in N2B-containing NMDARs. *Upper panels*, Whole-cell currents from HEK293 cells expressing human NMDAR subunits, either hGluN1/hGluN2A (**A**) or hGluN1/hGluN2B (**B**). Currents were elicited by a 2.5 s application of glutamate (1 mM) in the continuous presence of glycine (0.1 mM) (holding potential, −70 mV). *Lower panels*, control antibody B1 (IgG1, gray circles) or human-derived DNRAb G11 (green circles) were added 75 s after a baseline recording of 5 sweeps and were included in the bath throughout the remaining period. Current amplitudes for individual recordings were normalized to its baseline. Values are mean ± SEM (hN2A+B1, n = 6; hN2A+G11, n = 6; hN2B+B1, n = 5; hN2B+G11, n = 5). Example traces in upper panels show the +G11 recordings for the initial sweep during baseline (no antibody present) or for the last sweep during steady-state (in antibody). (**C & D**) Peak current amplitudes in N2A-containing NMDAR are more strongly potentiated than those in N2B-containing receptors. Bar graphs (mean ± SEM with dots indicating individual values) (from left to right for hN1/hN2A, n = 6, 6, 6, 6, 5, 5; and for hN1/hN2B, n = 5, 5, 6, 5) showing normalized steady-state peak current amplitudes either for control antibody (B1) or DNRAbs (G11). Significance of DNRAb values are measured relative to their respective control (\**p < 0.05* or \*\**p < 0.01, t-test*). *nt*, not tested. (**E & F**) Example single channel recordings of hGluN1/hGluN2A (**E**) and hGluN1/hGluN2B (**F**) with B1 (left traces) or G11 (right traces) antibodies at 10 μg/ml. Recordings were made in the on-cell configuration (holding potential, +100 mV). Downward deflections are inward currents. Low resolution top traces show 12 s of recording (filtered at 1 kHz); more high resolution bottom traces (selection in black box) show 230 ms (filtered at 3 kHz). (**G**) Equilibrium open probability (Eq. P_open_) (mean ± SEM) for N2A- or N2B-containing NMDARs (from left to right for hN1/nN2A, n = 6, 7, 8, 8; for hN1/hN2B, n = 6, 6, 6, 10). Antibody was either B1 (gray) or G11 (green) (\**p < 0.05, t-test*). (**H**) DNRAbs enhance forward rates of activation. Frequency of single channel openings (mean ± SEM) with antibody either B1 (gray) or G11 (green) (\**p < 0.05, t-test*). Test DNRAbs was 100 μg/ml.

For N2A-containing receptors, current amplitudes showed significant potentiation even at 1 μg/ml compared to its matched control (Figure 1C). In contrast, only at 100 μg/ml, did N2B-containing receptors show significant potentiation relative to its control (Figure 1D). Overall, these results suggest that, in terms of receptor gating, N2A-containing receptors are about 100-fold more sensitive to DNRAbs than N2B-containing receptors.

DNRAbs might potentiate NMDAR activity by acting as an agonist at the glutamate ligand-binding domain. To test this idea, we measured leak currents in the absence of glutamate in parallel to peak current amplitudes (Supplemental Table 1), but found that changes in leak current over time were not different in the presence of DNRAbs and their matched control. Hence, DNRAbs by themselves do not act as NMDAR agonists.

### DNRAbs act as positive allosteric modulators (PAMs)

To further verify these actions of DNRAbs, we measured single channel activity of N2A- or N2B-containing NMDARs exposed either to control antibody B1 or to G11 (Figures 1E-1H). These recording were made in the on-cell configuration in the continual presence of glutamate and glycine (as well as 0.05 mM EDTA)^24^. The equilibrium open probability (eq. P_open_), an index of the ease of ion channel opening, with control antibody for wild type hN1/hN2A (0.40 ± 0.07, n = 6)(mean ± SEM, n = number of patches) and hN1/hN2B (0.18 ± 0.04, n = 6) is comparable to previously published values^25^. At 10 μg/ml of G11, hN1/hN2A single channel activity was significantly potentiated (Figure 1E) relative to the matched control (Figure 1G). For hN1/hN2B, we again saw no significant effect of G11 on receptor gating at 10 μg/ml (Figures 1F & 1G). At 100 μg/ml (Figure 1G), the weak potentiation was not significant reflecting in part the variability of single channel recordings of N2B-containing receptors. These results further indicate that N2A-containing receptors are more sensitive to DNRAbs than N2B-containing receptors.

To define how DNRAbs potentiate activity, we characterized single channel details. Mean open (MOT) time were not significantly altered for either N2A- or N2B-containing receptors, whereas mean closed time (MCT) was significantly reduced for N2A-containing receptors at the highest concentration tested (Supplemental Table 2). Further, the frequency of opening was significantly enhanced for N2A-containing receptors (Figure 1H). Thus, DNRAbs act by enhancing forward rates to the open state rather than by stabilizing the open channel.

### A single copy of GluN2A confers high DNRAb sensitivity to NMDARs

At native synapses, NMDARs are typically composed of GluN1 in combination with different GluN2 subunits, often one GluN2A and one GluN2B subunit, so-called triheteromeric receptors^26, 27^. We therefore tested whole-cell currents from triheteromeric receptors, where the receptor contains a single copy of GluN2A (Figure 2A). To generate triheteromeric receptors, we used NMDAR constructs having coiled-coiled domains that permit only specific subunit combinations to reach the plasma membrane^28^. At 10 μg/ml of DNRAb, diheteromeric N2A-containing receptors containing the coiled-coiled domains showed significant potentiation. Diheteromeric N2B-containing receptors again showed no potentiation. In contrast, triheteromeric receptors containing a single copy of GluN2A showed significant potentiation. Thus, a single copy of GluN2A confers high sensitivity to DNRAbs.

**Figure 2.**
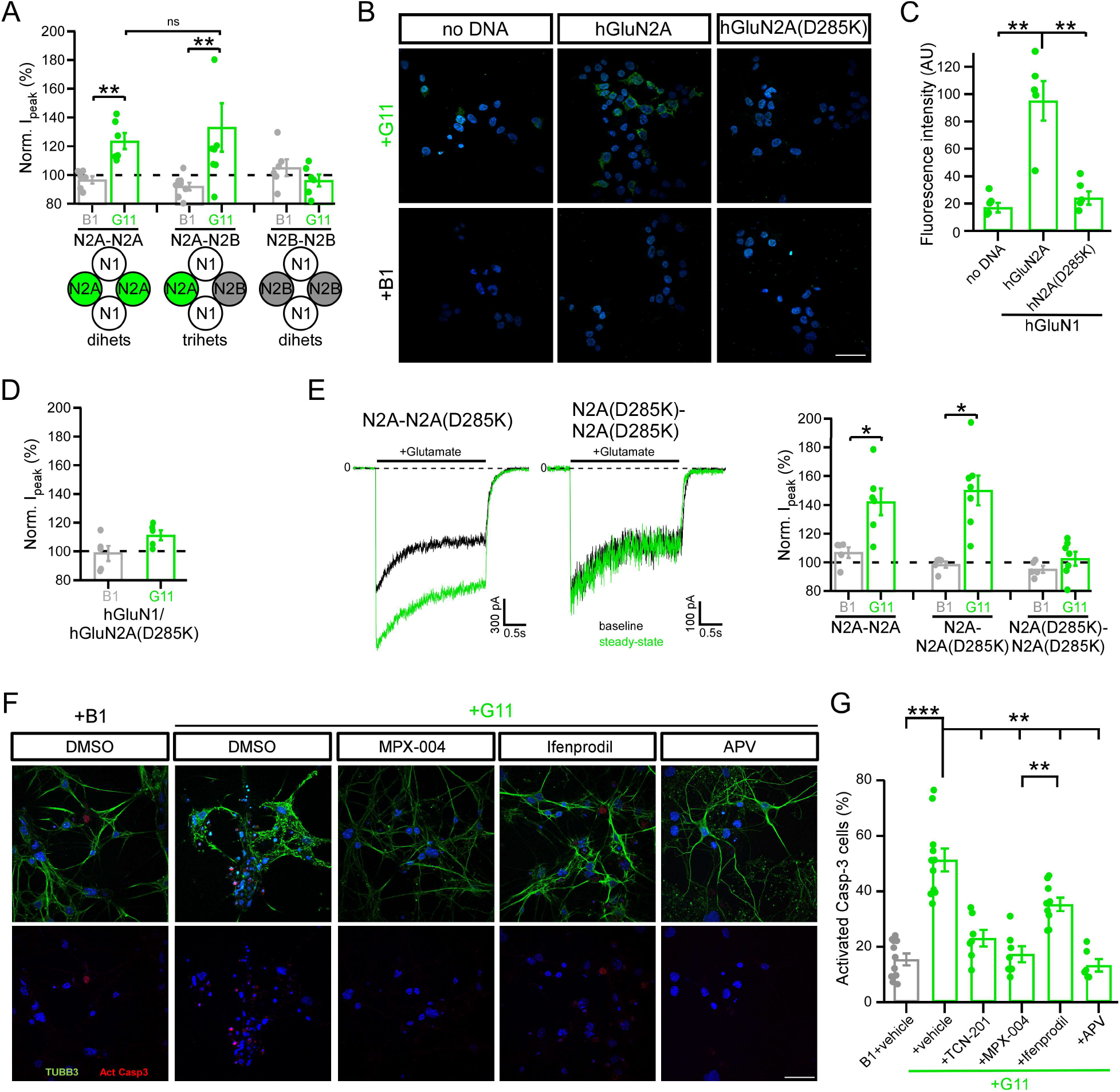
A single copy of GluN2A confers high sensitivity to NPSLE DNRAbs. **(A)** Bar graphs (mean ± SEM) (from left to right, n = 6, 6, 8, 8, 6, 6) showing normalized whole-cell peak current amplitudes for diheteromeric NMDARs (N1/N2A-N2A or N1/N2B-N2B) or triheteromeric receptors (N1/N2A-N2B) at 10 μg/mL of B1 or G11 antibody (\*\**p < 0.01, Mann-Whitney U test*). **(B)** A single charge reversal in the GluN2A DWEYS motif (DW<u>D</u>YS) (D285K) disrupts G11 binding. Representative images from immunocytochemistry of HEK293T cells not transfected (no DNA) or transfected with GluN1/GluN2A or with GluN1/GluN2A(D285K). Cells were stained with either B1 or G11 (10 μg/mL) (green, Alexa-488) under non-permeabilizing conditions with DAPI (blue) counterstain. Scale (white bar): 40 μm. **(C)** Quantification of mean fluorescence intensity in (B) (mean ± SEM) (n = 5, all conditions) (\*\**p < 0.01, one-way ANOVA with post-hoc Tukey’s test*). **(D)** Single charge reversal prevents DNRAb (10 μg/mL) potentiation of whole cell currents (mean ± SEM, n = 5, all conditions). **(E)** *Left*, Current records of triheteromeric (N1/N2A-N2A(D285K)) or diheteromeric (N1/N2A(D285K)-N2A(D285K)) NMDARs. Currents displayed as in Figure 1A. *Right*, Bar graphs (mean ± SEM) (from left to right, n = 5, 6, 5, 7, 5, 6) showing normalized current amplitude for diheteromeric NMDARs (N1/N2A-N2A or N1/N2A(D285K)-N2A(D285K)) or triheteromeric receptors (N1/N2A-N2A(D285K)) at 10 μg/mL of B1 or G11 (\**p < 0.05, Mann-Whitney U test*). **(F)** GluN2A specific antagonists are neuro-protective at moderate pathophysiological levels of DNRAbs. Immunocytochemistry of primary hippocampal neuronal cultures (DIV14) incubated either in control antibody (B1+vehicle) or in DNRAb (G11) at 10 μg/mL. G11 was incubated either alone (+vehicle), with GluN2A antagonists TCN-201 (not shown) or MPX-004, or with ifenprodil, a GluN2B negative allosteric modulator at 3 μM. Upper panels, representative images of hippocampal neurons stained with antibodies against neuronal-specific β-tubulin, Tubb3 (green, Alexa-488), and activated caspase-3 (red, Alexa-647), with DAPI (blue) counterstain. Lower panel, only activated caspase-3 channel is shown. Scale (white bar): 40 μm. **(G)** Quantification of immunocytochemistry results shown in (F). Proportion of DAPI and activated-caspase-3-positive cells divided by total DAPI cells (mean ± SEM) (from left to right, n = 11, 11, 7, 7, 9, 6) per treatment condition (\**p < 0.05, **p < 0.01, ***p < 0.001, one-way ANOVA with post-hoc Tukey’s test*).

### DNRAbs modify receptor function via the DWEYS motif

DNRAb interact and presumably alter NMDAR function through the DWEYS motif in the amino-terminal domain. To directly test this idea, we introduced a charge reversal in the middle of the GluN2A DWEYS motif (DWDYS), mutating the negatively charged aspartate (D) at position 285 (DW<u>D</u>YS) to the positively charged lysine (K) (D285K). We avoided sites possibly involved in coordinating Zn^2+^ to mitigate confounding effects from Zn^2+^ modulation^29, 30^. N2A(D285K) has no apparent effect on current properties (Supplemental Table 3). The G11 antibody shows robust binding to NMDARs (Figure 2B, middle images). This binding is significantly attenuated in receptors containing D285K (Figures 2B, right images, & 2C). We did not find significant differences in the intrinsic binding of GluN2A or GluN2B-specific epitopes to G11 (Supplemental Figure 1), suggesting that the difference in sensitivity primarily reflects gating. Receptors containing D285K are no longer significantly potentiated by DNRAbs (Figure 2D), indicating that binding of DNRAbs to the DWEYS motif mediates the positive allosteric effect.

We also tested whether a single (triheteromeric receptor) or two (diheteromeric receptors) copies of the charge reversal are required to disrupt function. Consistent with the results for GluN2A/GluN2B triheteromeric receptors, a single copy of wild type GluN2A confers full sensitivity to DNRAbs (Figure 2E). Thus, in terms of stoichiometry, binding of a single DNRAbs to the DWEYS motif on GluN2A can generate the full positive allosteric effect.

### GluN2A antagonists are neuroprotective against DNRAbs

Our results indicate that N2A-containing receptors are significantly more sensitive to DNRAbs than N2B-containing receptors. To see if attenuating N2A-activity is neuroprotective, we incubated primary hippocampal cultures either in control antibody (B1) (Figure 2F, left images) or in G11 (10 μg/mL) in the absence or presence of N2A antagonists (TCN-201 or MPX-004), which are specific at the concentrations used here^28, 31^, or a N2B-specific negative allosteric modulator (ifenprodil) and assayed the number of DAPI positive cells associated with activated caspase-3 activity (Figure 2G). In the presence of DNRAb alone, apoptotic cell death was extensive, consistent with previous results^3, 8, 32^. This cell death was significantly attenuated in the presence of MPX-004. In contrast, ifenoprodil was less able to protect against cell death (Figures 2F & 2G). Thus, specific N2A-antagonists are neuroprotective against DNRAb-mediated cell death.

### Assaying the contribution of GluN2A and GluN2B NMDAR subunits to DNRAb-induced phenotype *in vivo*

To address how the different NMDAR subunits contribute to the DNRAb-induced phenotypes *in vivo*, we used transgenic mouse models, either with the GluN2A subunit knocked out (*grin2A^−/−^* mice, termed ‘N2A KO’ henceforth) or with a conditional knockout of the GluN2B subunit (*grin2B*^fl/fl^; *CaMKII*^Cre^ mice, termed ‘N2B cKO’) (Figure 3A). We used the conditional KO due to the embryonic lethality of a full GluN2B knockout^33, 34, 35^. KO mice were immunized with a decapeptide containing the pentapeptide, DWEYS, a mimetope of dsDNA and homologous to a sequence within the GluN2A and GluN2B extracellular domains, multimerized on a polylysine backbone (MAP-DWEYS) (Supplemental Figure 2). Immunization of wild type mice with MAP-DWEYS induces production of DNRAbs (DNRAb+ mice)^2, 11, 36^. As a control, mice were also immunized with the polylysine backbone alone (DNRAb– mice). Two weeks following two booster immunizations, mice were given LPS to allow transient access of antibodies to the hippocampus (Supplemental Figure 2)^11, 19^.

**Figure 3.**
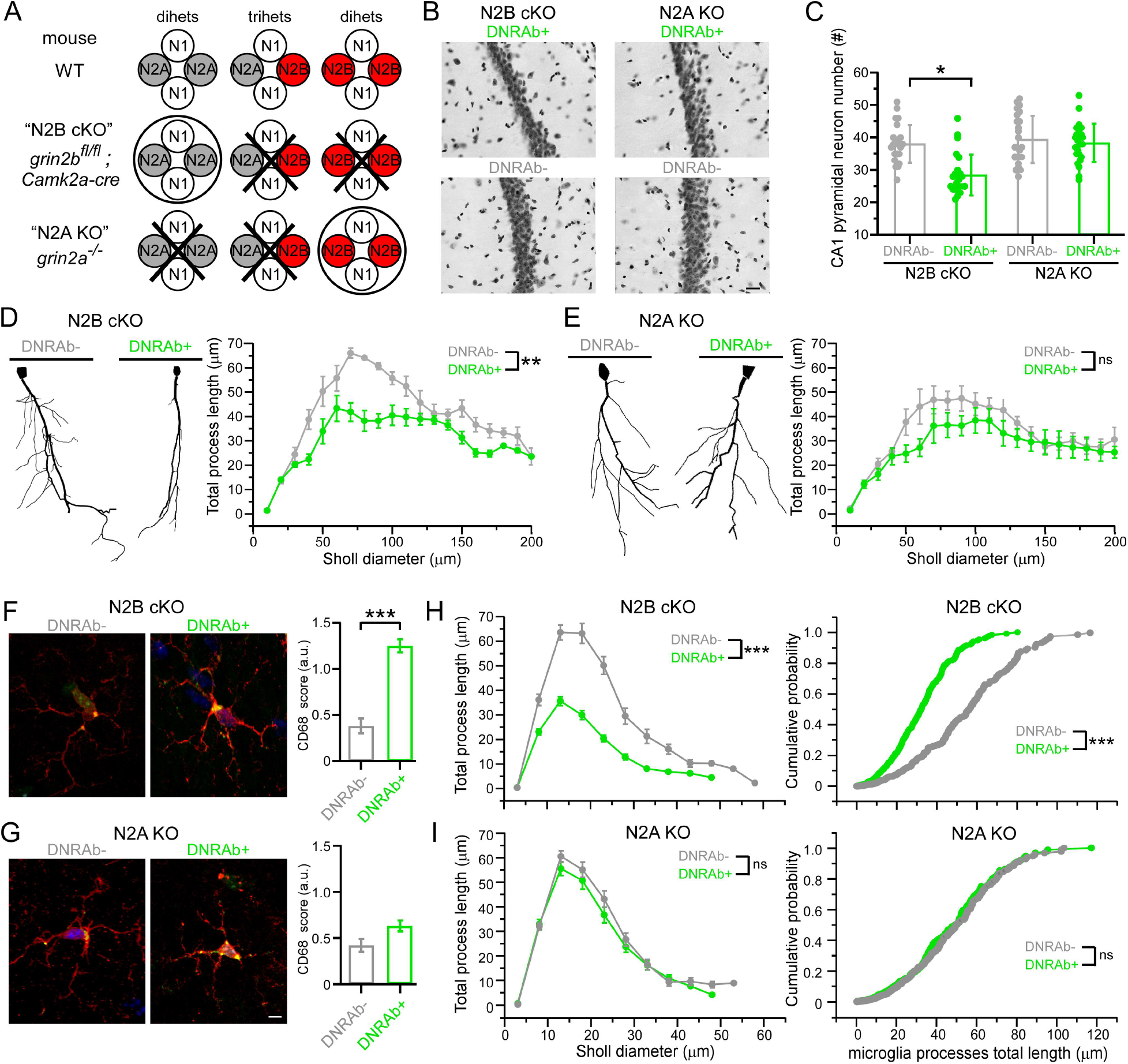
DNRAbs induced neuronal loss and dendritic pruning in CA1 pyramidal neurons persist in mice lacking GluN2B, but not in those lacking GluN2A. **(A)** Schematic of NMDAR composition expected in hippocampal pyramidal neurons in the various genetic backgrounds. N2B cKO and N2A KO mice were immunized with either MAP-core (DNRAb-) or MAP-DWEYS (DNRAb+), and subsequent to LPS treatment, were sacrificed and their hippocampi assayed for cell death (1 week post LPS) or cellular morphology (8 weeks post LPS) (Supplemental Figure 2). **(B)** Micrographs of hippocampal CA1 neurons from DNRAb+ or DNRAb- N2B cKO or N2A KO mice. Scale bar, 20 μm. **(C)** Quantification of CA1 neurons (mean ± SEM, n=3). Each dot represents a CA1 pyramidal neuron counted using a standard unbiased stereological protocol (8 runs of systematic random sampling in 3 animals for each group). (\**p<0.05, Mann-Whitney U test*). **(D & E)** DNRAb+ N2B cKO mice show significantly reduced dendritic complexity. Left, Tracings of Golgi-stained CA1 pyramidal neurons from N2B cKO (**D**) or N2A KO (**E**) mice. Right, Sholl analysis, which measures dendritic complexity, for N2B cKO (**D**) or N2A KO (**E**) mice (** p < 0.01, Kolmogorov-Smirnov test). Values at concentric rings are shown as mean ± SEM. For each group, 8 runs of systematic random sampling was done from three (3) animals. **(F & G)** Representative images of microglia from N2B cKO (A) or N2A KO (B) mice. Microglia are labeled with antibody to Iba1 (red, Alexa 594) and to CD68 (green, Alexa 488)]. Scale (white bar): 20 μm. Right panels, quantification of CD68+ score (see Supplemental Methods) (*** *p* <*0.001, Kruskal-Wallis ANOVA test*). **(H & I)** Microglia in DNRAb+ N2B cKO mice show significant diminution of complexity and process length. Left panels, Sholl analysis of process complexity of microglia from N2B cKO (**H**) or N2A KO (**I**) mice (\*\*\**p< 0.001, Kolmogorov-Smirnov test*). *Right panels*, cumulative probability curves of microglia process length from N2B cKO (**H**) or N2A KO (**I**) (\*\*\**p< 0.001, Kolmogorov-Smirnov test*). Three (3) animal were used per group; number of quantified microglia: N2B cKO, DNRAb- (n = 33), DNRAb+ (n = 34); N2A KO, DNRAb− (n = 39), DNRAb+ (n = 40).

Wild type mice, when immunized with MAP-DWEYS, show two stages of DNRAb-induced pathology (Supplemental Figure 2)^2, 11, 19^. An acute phase, assayed one week after LPS treatment, that is characterized by extensive cell death of hippocampal pyramidal neurons^11^; and a chronic phase, assayed eight weeks post-LPS treatment, characterized by the loss of dendritic complexity in pyramidal neurons, microglia activation, and disruption of place cell function (Supplemental Figure 2). DNRAb+ mice also display reduced spatial memory, presumably mimicking the phenotype in SLE patients^2, 11^. Because LPS only transiently permeabilizes the blood brain barrier^2^, the acute phase is associated with the presence of DNRAbs, whereas the chronic phase is independent of the direct presence of DNRAbs^2, 19^.

### Mice lacking GluN2B show DNRAb-mediated pathology while mice lacking GluN2A are protected

Initially, we assayed the effect of DNRAbs on the acute phase of the pathology, namely the induction of pyramidal neuron cell death measured one week after LPS treatment^11, 19^. For all *in vivo* experiments, we compare the effect in DNRAb+ mice to that in DNRAb- mice in the same genetic background. DNRAb+ mice in the N2B cKO background, which still retain GluN2A, also displayed a significant loss of hippocampal neurons (Figures 3B & 3C). In contrast, N2A KO mice showed no significant loss of neurons. Hence, the loss of pyramidal neurons during the acute phase is dependent on GluN2A.

Next, we characterized the chronic phase of pathology by assaying pyramidal neuron morphology eight weeks after LPS administration (Figures 3D-3I). Compared to its control, GluN2B cKO mice exposed to DNRAbs exhibited a significant loss of dendritic complexity, as assayed by Sholl analysis (Figures 3D). In contrast, in GluN2A KO mice, DNRAbs have no significant effect on dendritic complexity (Figure 3E). Thus, as with acute enhanced cell death, the chronic phase of SLE pathology associated with reduced dendritic complexity is also dependent on GluN2A.

### DNRAb-induced activation of microglia does not occur when GluN2A is absent

The reduced dendritic complexity occurring during the chronic phase of DNRAb-induced pathology is dependent on microglia activation^19^. We therefore characterized microglia in DNRAb+ and DNRAb- mice in the various genetic backgrounds (Figures 3F-3I).

For the N2B cKO background, and in comparison to its control, DNRAb+ mice showed enhanced expression of CD68 (Figure 3F, right panel) and diminution of process complexity (Figure 3H, left panel), and process length (Figure 3H, right panel) over concentric Sholl diameters. These results are consistent with microglia activation and parallel what is observed in wild type mice^19^. In contrast, for the N2A KO background (Figures 3G & 3I), DNRAb+ mice, in comparison to its control, did not show significant CD68 expression (Figure 3G, right panel) or reduced complexity (Figure 3I), suggesting a lack or reduced microglia activation. Thus, DNRAb-induced microglia activation, which is critical in dendritic morphology changes, either directly requires GluN2A and/or is dependent on acute GluN2A-induced cell death.

### GluN2A is required for DNRAb-induced changes in spatial memory

During the chronic phase of DNRAb-induced pathology, wild type mice exposed to DNRAb show impaired spatial memory and disrupted place cell properties in the CA1 region of the hippocampus (Supplemental Figure 2)^2, 11, 19^. We therefore asked whether DNRAb+ N2A KO and N2B cKO mice, 8 weeks after LPS exposure, behaved abnormally in an object-place memory (OPM) task, which tests spatial memory (Figure 4A)^37, 38^. N2B cKO mice transiently exposed to DNRAbs performed significantly worse than its DNRAb- control in the OPM task (Figure 4B). This parallels to what is observed in wild type mice (Supplemental Figure 2). In contrast, no difference in performance was detected between DNRAb+ and DNRAb– N2A KO mice. Thus, GluN2A, either directly or indirectly, is required for DNRAb-induced disruptions in spatial memory.

**Figure 4.**
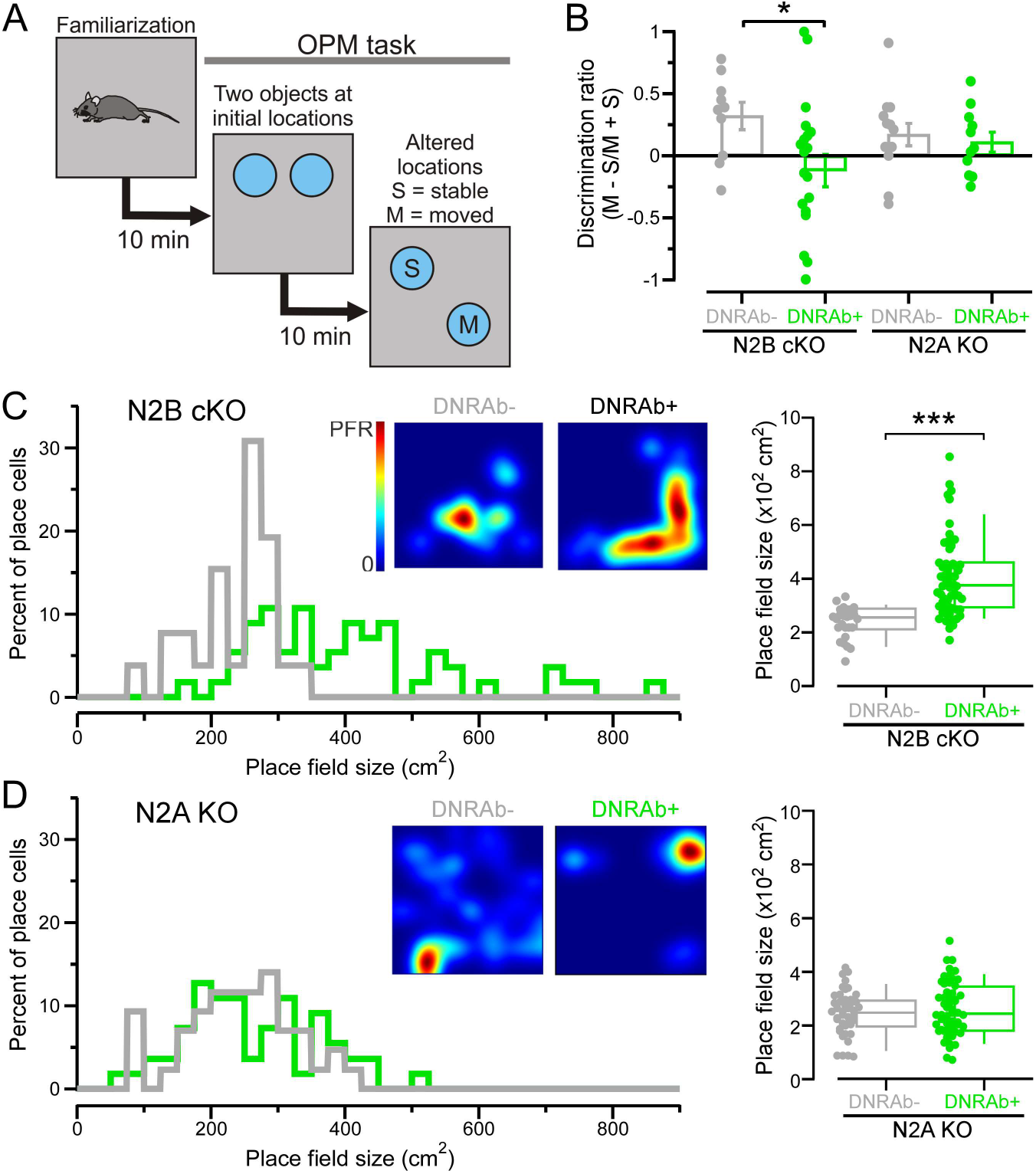
Behaviors disrupted by DNRAbs persist in mice lacking GluN2B, but not in those lacking GluN2A. **(A)** Schematic of the object place memory (OPM) task. **(B)** DNRAb+ N2B cKO mice show significantly reduced exploratory behavior, relative to its control, of the moved (novel) object over the stable object (Supplemental Figure 2). Mice were tested 8 weeks after LPS treatment. Values shown are mean ± SEM (n = 10, 20, 13, 11, from left to right, number of tested mice per group; \**p < 0.05, Mann-Whitney U test*). **(C & D)** DNRAb disrupts place field size in mice lacking GluN2B, but not in those lacking GluN2A. *Left*, Histograms for place field sizes of all place cells recorded in DNRAb- (gray) or DNRAb+ (green) mice either N2B cKO (**C**) or N2A KO (**D**) backgrounds. *Inset*, Heatmaps of representative place fields. *Right*, Box plots (25^th^ & 75^th^ percentiles with the whiskers, 10 and 90 percentile), comparing place field size in N2B cKO (**C**) or N2A KO (**D**) mice (*** *p* < *0.001, Mann-Whitney U test*). Each dot represents a single place field: for N2B cKO (**C**), DNRAb− (n = 26) and DNRAb+ (n = 56); for N2A KO (**D**), DNRAb− (n = 43) and DNRAb+ (n = 55).

NMDAR-dependent spatial memory and learning is linked to hippocampal place field size^39, 40^. In wild type mice exposed to DNRAbs, place field size is aberrantly enlarged, paralleling the reduced spatial memory in SLE patients and other neurodegenerative models^2, 19, 41^. To assess neuronal functional integrity, we measured place field size in freely-moving mice by implanting tetrodes into the dorsal CA1 region (Figures 4C-4D). As in wild type, DNRAb+ N2B cKO mice exhibited significantly larger place field sizes compared to DNRAb– N2B cKO mice (Figure 4C), indicating that in the absence of GluN2B, the DNRAb-induced disruption of place field size persists. In contrast, DNRAb+ and DNRAb– N2A KO mice displayed no significant differences in place field size (Figure 4D). Thus, the cognitive deficits associated with DNRAbs are intimately linked to GluN2A, but appear largely independent of GluN2B.

## DISCUSSION

Our experiments demonstrate that the GluN2A NMDAR subunit is required for various neuropathologies associated with lupus auto-antibodies. We find that DNRAbs act as positive allosteric modulators (PAM) on NMDARs with the GluN2A subunit showing a much higher sensitivity than the GluN2B subunit (Figure 1). DNRAb exposure manifests in two distinct phases: First, synaptic disturbances and cell death with acute exposure, and second, a loss of dendritic complexity resulting from microglia activation that occurs even after antibody is no longer detected^10, 19^. Presumably, the higher sensitivity of GluN2A to DNRAbs facilitates cell death during acute exposure. The chronic phase, including the cognitive and memory defects similar to what is seen in SLE patients^2, 42^, are also dependent on GluN2A but it is unknown whether it is a residual consequence of excitotoxicity encountered during the acute phase or a lasting change to NMDARs that predispose neurons to microglial-dependent pruning.

G11, the human DNRAb we tested, acts as a PAM on NMDARs via the DWEYS-motif (Figures 2B-2D). DNRAbs are present in about 30-40% of SLE patients, with these DNRAbs being identified by binding to peptides containing the DWEYS-motif^9, 10, 14, 43, 44^. Hence, we assume that the PAM action is a common feature of all DNRAbs. Consistent with this idea, polyclonal antibodies from the CSF of SLE patient with neuropsychiatric symptoms that bind the DWEYS-motif induce NMDAR hyperfunction^12, 32^. Nevertheless, it is possible that there are clonal variations in the magnitude of the PAM action as well as possible additional functional effects of DNRAbs, necessitating the study of more SLE patient DNRAbs. The concentration of DNRABs in the nervous system presumably varies during the pathophysiological course of SLE. At low concentrations, DNRAbs would affect exclusively GluN2A-containing receptors, while at high concentrations, DNRAbs would also affect GluN2B diheteromeric receptors (Figure 1). Previous work has shown that the GluN2B-specific inhibitor, ifenprodil, can be neuroprotective^10^. However, this most likely reflects that tri-heteromeric GluN1/GluN2A/GluN2B NMDARs, which constitute a major portion of hippocampal synapses^26, 27^, are inhibited by ifenprodil^28^. Indeed, we find that a single copy of wild type GluN2A confers high sensitivity to DNRAbs (Figure 2A), a result consistent with GluN2A dominance in allosteric modulation^45, 46^.

The DWEYS-motif is located in the amino-terminal domain (ATD) clamshell hinge region (Figures 2B-2D). GluN2A-selective PAMs and negative allosteric modulators (NAMs) have been developed to target NMDAR hypo- or hyperfunction in disease^47, 48^. While most GluN2A-selective PAMs and NAMs target the ligand-binding domain (LBD), the ATD interacts with the LBD to mediate its allosteric action^49, 50, 51, 52^. In GluN2A/GluN2B NMDARs, the GluN2A ATD interacts more extensively with the LBD than the GluN2B ATD^45, 46^, which may underlie the higher sensitivity of GluN2A to DNRAbs. Furthermore, the DWEYS motif is proximal to binding sites for other ATD allosteric modulators. Indeed, the sensitivity to Zn^2+^, a NAM, may be modulated by SLE antibodies^32^. Nevertheless, the mechanism of action of DNRAbs to produce positive allosteric modulation and the differences in sensitivity between GluN2A and GluN2B remain unknown, but are critical to define for potential therapeutic interventions.

GluN2A contributes to both the acute and chronic phases of SLE neuropathology. While the use of GluN2A NAMs may be key to treating the neuropsychiatric symptoms of SLE, any NMDAR inhibitor may lead to undesirable side effects^20, 53, 54^. Uncovering the mechanistic details of DNRAb-NMDAR interactions during distinct pathophysiological phases will help us better develop and tailor therapeutic strategies for preventing neuropsychiatric symptoms in SLE patients before acute events occurs, and treating them during chronic phases of SLE.

## METHODS

See Supplemental Methods for details.

### Animals

#### Feinstein Institute

Mice with deletion of the GluN2A subunit (*Grin2a*^−/−^) were kindly provided by Dr. M. Mishina (University of Tokyo). Mice with conditional knockout of the GluN2B subunit (*Grin2b*^fl/fl^;*Camk2a-cre*) were bred in house by crossing homozygous floxed GluN2B mice (*Grin2b*^fl/fl^, kind gift from Prof. Dr. H. Monyer) with a Cre line driven by CaMKIIα promoter (*Camk2a-cre* mice, B6.Cg-Tg(Camk2a-Cre)T29-1Stl/J, stock no: 005359, Jackson labs). Both groups were further bred on an H2d^+/+^ background to allow antibody response to immunization. Immunization with MAP-core and MAP-DWEYS was previously described^11^.

#### Stony Brook

Mice (C57BL/6) were bred in house for primary hippocampal cultures at P0-P1 and glia/astrocyte feeder layers at P2-P4.

### SLE antibodies

G11 is an IgG1 monoclonal antibody derived from a female SLE patient’s B cells with Igγ1 heavy chain and Igκ light chain composition. B1 is an isotype IgG1 control derived from the same patient^16^. G11 was cloned from a B cell binding the DWEYS peptide and B1 from a B cell that did not bind the peptide or any brain antigen. The SLE patient from whom the antibodies were derived had serum antibodies reactive to dsDNA and the DWEYS peptide^2, 4^.

### *In vitro* electrophysiology

Human embryonic kidney 293 (HEK293) cells were grown and transfected (Supplemental Methods: *in vitro* cell culture and transfection). Whole-cell^55, 56^ and single-channel recordings^24, 57^ were done as described (Supplemental Methods: *in vitro* macroscopic recordings and *in vitro* single channel recordings).

### Immunocytochemistry (ICC)

HEK293T cells were used for ICC. Cells were fixed 48 hours post-transfection, washed with 1xPBS, and then blocked with 2% bovine serum albumin (BSA) (Sigma, A9647). Incubation of primary antibody (20 μg/mL), either G11 or control B1, diluted in 2% BSA was done overnight at 4°C.

Primary hippocampal neurons were made^58^ with minor modifications (see Supplemental Methods). Pharmacological treatment occurred at DIV14. Final concentrations of antagonists were 3 μM: TCN-201 (AdooQ Bioscience, A11947), MPX-004 (Alomone Labs, M280), and Ifenprodil (Sigma, I2892). ICC was performed 24-hours post treatment on DIV15. Coverslips were washed with 1xPBS, fixed, and washed again with 1xPBS and blocked/permeabilized with antibody blocking solution, consisting of 2% (w/v) BSA, 0.25% (v/v) Triton-X100 in 1xPBS, for 60 min. Coverslips were then incubated in primary antibodies, rabbit anti-activated caspase-3 (1:250, Cell Signaling, 9661), and mouse anti-β tubulin III (Tubb3) (1:400, Millipore-Sigma, MAB1637), both in antibody blocking solution overnight in 4°C. Coverslips were washed with 1xPBS, and incubated in fluorescence-conjugated secondary antibodies, Goat anti-rabbit Alexa-647 (1:1000, ThermoFisher, A21245) and Goat anti-mouse Alexa-488 (1:1000, ThermoFisher, A21121) for 1 hour at RT and mounted as described above. Post-processing and image analysis are detailed in Supplemental Methods.

### Immunohistochemistry

Mice were anesthetized as described^19^. Brains were extracted, fixed, and transferred to 30% sucrose, blocked and sliced (40 μm). For cresyl violet staining, sections were mounted, dehydrated, rehydrated and stained in cresyl violet. Sections were dried, dehydrated, cleared (Histoclear II) and coverslipped (Permount, Fisher Scientific) prior to imaging on an AxiophotZ1 microscope (Zeiss). For immunohistochemistry, sections were washed with PBS, permeabilized, blocked, and stained with primary antibody overnight at 4°C. Primary antibodies were Iba1 (1:500, Wako Chemicals, 019-1 9741), CD68 (1:500, Bio-Rad, mca1957). On the following day, samples were washed with PBS and then incubated with secondary antibody, which included Donkey anti-rat Alexa Fluor 488 (1:400, Life Technologies, A21208), Chicken anti-rabbit Alexa Fluor 594 (1:300, Life Technologies, A21442). Post-processing and image analysis are detailed in Supplemental Methods.

### Behavioral assessment and *in vivo* electrophysiology

Mice (20-24 weeks) were handled for 15 min per day for 3 days before being tested behaviorally. The object place memory (OPM) task was performed^2, 19^. The apparatus consisted of a square (40 cm) chamber 40 cm high, with the walls painted grey. Animals were familiarized to the empty chamber (3 sessions of 15 min each). For OPM testing, mice underwent the following sequence: empty chamber (10 min), home cage (10 min), sample phase in which the chamber had two objects located at the center of the NW and NE sectors (5 min), home cage (10 min), choice phase in which the chamber had the same objects but one of them was moved from NE to the center of the SE sector (5 min). The discrimination ratio was calculated during the choice phase by dividing time spent exploring the moved object minus the time spent exploring the static object by the time exploring the objects combined^2^. Data were collected and analyzed using EthoVision XT (Noldus Information Technologies).

*In vivo* recordings of place cells in the dorsal CA1 of the hippocampus were made^2, 19^. Mice were anesthetized and implanted with a 16 channel multi-electrode array. After recovery, single unit firing was recorded using Cheetah software (Neuralynx) while the animal explored a square chamber in a schedule of four exploration runs separated by three rest sessions. Data were analyzed using Spike2 (version 8, Cambridge Electronic Design), NeuroExplorer (version 5, Nex Technologies), and Matlab.

### Statistics

Data analysis was performed using IgorPro, QuB, Excel, and ImageJ. All average values are presented as mean ± SEM. For statistical analysis, we used either Origin Pro (version 9, Origin Lab) or MiniTab 18. Normality was used to determine appropriate statistical tests. For normally distributed data, unpaired two-tailed Student’s t-tests (*t-test*) or one-way ANOVAs with post-hoc Tukey’s test were used as indicated. For non-normally distributed data or instances where the data were categorical variables or cumulative distribution functions, Mann-Whitney U tests, Kruskal-Wallis ANOVAs, and Kolmogorov-Smirnov tests were used as indicated. Significance was typically *p* < 0.05.

We did not run a statistical test to determine sample size *a priori*. Sample sizes resemble those from previous publications.

## Acknowledgments

We thank Donna Schmidt for technical assistance; Dr. Quan Gan for helpful discussions and/or comments on the manuscript; Drs. Stephen Traynelis and Hongjie Yuan (Emory University) for generously sharing human NMDAR constructs; Dr. Kasper Hansen (University of Montana) for supplying tri-heteromeric constructs; and Prof. Masayoshi Mishina (University of Tokyo) and Prof. Dr. Hannah Monyer (Universität Heidelberg) for sharing transgenic mice targeting NMDAR subunits. This work was supported by NIH Grants NRSA F30MH115618 (KC), R01 NS088479 (LPW), and P01A1073693 (BD).

## Author Contributions

K.C., J.N., P.T.H., B.T.V., B.D., and L.P.W. designed the experiments. K.C. J.N, T.S.H., N.C., G.M., C.K., and P.T.H. performed the experiments. K.C., J.N. P.T.H., B.T.V., B.D., and L.P.W. analyzed the data. K.C., P.T.H., B.T.V., B.D., and L.P.W. wrote the paper.

## SUPPLEMENTAL MATERIAL

**Supplemental Table 1.**
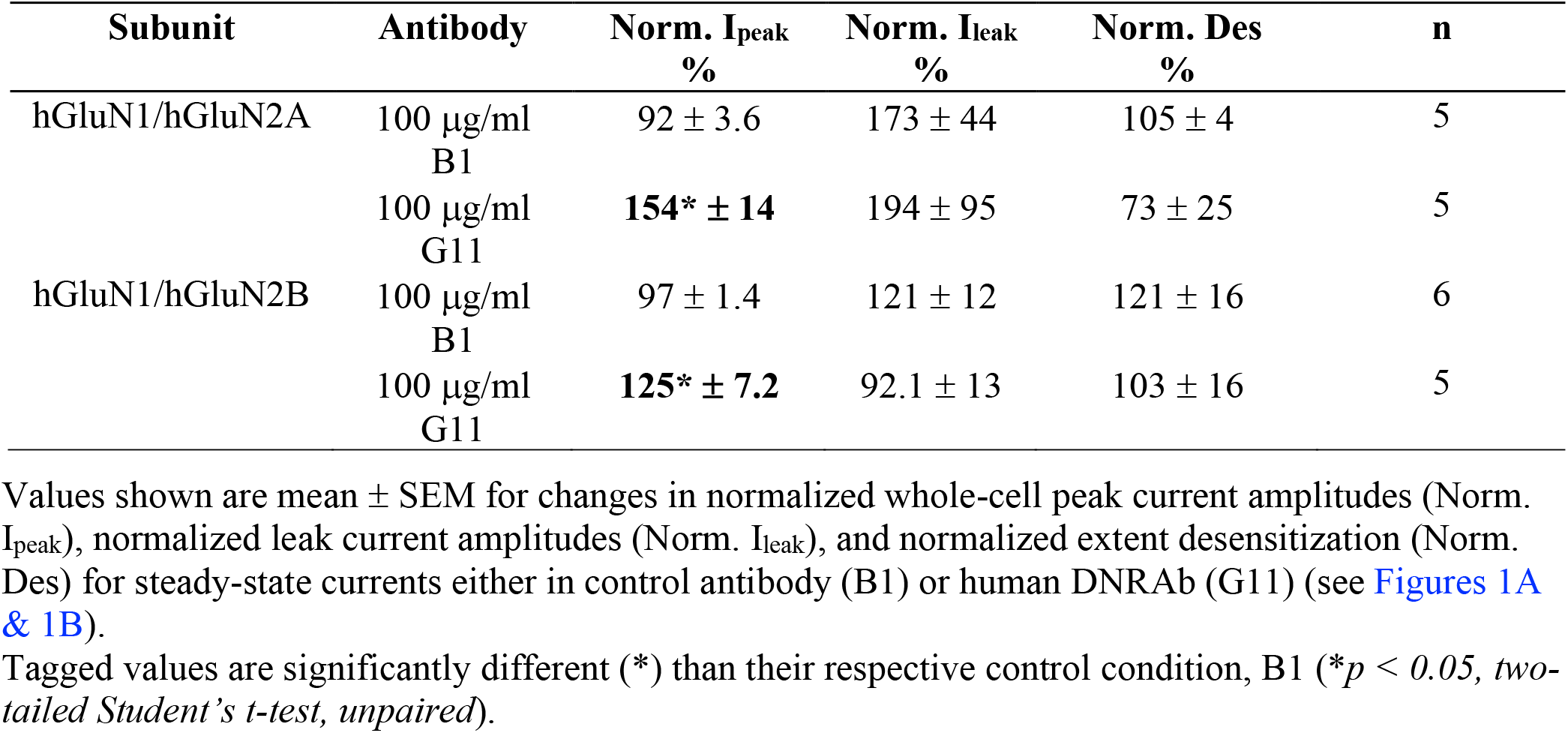
In the whole-cell mode, the G11 antibody has no notable effect on current properties other than changes in peak current amplitudes (relates to Figures 1A-1D).

| Subunit | Antibody | Norm. $I_{peak}$<br>% | Norm. $I_{leak}$<br>% | Norm. Des<br>% | n |
| --- | --- | --- | --- | --- | --- |
| hGluN1/hGluN2A | 100 μg/ml<br>B1 | $92 \pm 3.6$ | $173 \pm 44$ | $105 \pm 4$ | 5 |
| | 100 μg/ml<br>G11 | **$154^* \pm 14$** | $194 \pm 95$ | $73 \pm 25$ | 5 |
| hGluN1/hGluN2B | 100 μg/ml<br>B1 | $97 \pm 1.4$ | $121 \pm 12$ | $121 \pm 16$ | 6 |
| | 100 μg/ml<br>G11 | **$125^* \pm 7.2$** | $92.1 \pm 13$ | $103 \pm 16$ | 5 |

Tagged values are significantly different (\*) than their respective control condition, B1 ( $*p < 0.05$ , two-tailed Student's *t*-test, unpaired).

We made these measurements in 100 μg/ml where both N2A- and N2B-containing receptors were potentiated (Figures 1C & 1D). The lack of an effect on leak current indicates that DNRAbs do not themselves act as agonists. DNRAbs also do not affect desensitization.

**Supplemental Table 2.** Effects of DNRAbs on hGluN2A- or hGluN2B-containing NMDARs (relates to Figures 1E-1H).

| Subunit | Antibody | Total events (# of patches) | $i$<br>$pA$ | eq. $P_{open}$ | MCT<br>$ms$ | MOT<br>$ms$ |
| --- | --- | --- | --- | --- | --- | --- |
| hGluN2A | 10 $\mu g/ml$ B1 | 485793 (6) | $-8.3 \pm 0.4$ | $0.40 \pm 0.07$ | $5.9 \pm 1.0$ | $3.8 \pm 0.6$ |
| | 10 $\mu g/ml$ G11 | 546303 (7) | $-8.2 \pm 0.5$ | **$0.63^* \pm 0.05$** | $3.5 \pm 0.7$ | $6.0 \pm 0.9$ |
| | 100 $\mu g/ml$ B1 | 632383 (8) | $-7.1 \pm 0.2$ | $0.40 \pm 0.07$ | $8.9 \pm 1.9$ | $4.9 \pm 0.7$ |
| | 100 $\mu g/ml$ G11 | 1047339 (8) | $-7.5 \pm 0.1$ | **$0.61^* \pm 0.03$** | **$3.7^* \pm 0.4$** | $5.7 \pm 0.4$ |
| hGluN2B | 10 $\mu g/ml$ B1 | 135134 (6) | $-8.3 \pm 0.3$ | $0.18 \pm 0.04$ | $27.9 \pm 12.4$ | $4.0 \pm 0.7$ |
| | 10 $\mu g/ml$ G11 | 445555 (6) | $-8.5 \pm 0.2$ | $0.19 \pm 0.02$ | $16.0 \pm 2.9$ | $3.5 \pm 0.3$ |
| | 100 $\mu g/ml$ B1 | 154900 (6) | $-7.5 \pm 0.4$ | $0.15 \pm 0.06$ | $42.9 \pm 12.9$ | $4.8 \pm 0.9$ |
| | 100 $\mu g/ml$ G11 | 332097 (10) | $-7.5 \pm 0.3$ | $0.23 \pm 0.04$ | $24.0 \pm 2.8$ | $6.1 \pm 0.5$ |

Values shown are mean ± SEM for single-channel current amplitude ( $i$ ), equilibrium open probability ( $P_o$ ), mean closed time (MCT), and mean open time (MOT). Single channel currents were recorded in the on-cell mode at approximately -100 mV and analyzed in QuB (see Materials & Methods). Number of patches is in parenthesis to the right of total events. Eq $P_o$ is the fractional occupancy of the open states in the entire single-channel recording, including long lived closed states (desensitized states). All data were idealized and fit at a dead time of 20 $\mu s$ .
Tagged values are significantly different (\*) than their respective isotype control antibody (Ab) condition, B1 ( $p < 0.05$ , two-tailed Student's $t$ -test, unpaired).

**Supplemental Table 3.** The D285K mutation in the GluN2A DWEYS motif has no notable effect on current properties (relates to Figures 2B-2D).

| Subunit | $I_{ampl}$<br>$pA$ | Des<br>% | n |
| --- | --- | --- | --- |
| hGluN1/hGluN2A | $-1200 \pm 550$ | $33 \pm 12$ | 10 |
| hGluN1/ hGluN2A(D285K)/hGluN2A(D285K) | $-970 \pm 290$ | $19 \pm 10$ | 7 |

Values shown are mean ± SEM for whole-cell peak current amplitude ( $I_{ampl}$ ) and % desensitization (Des). Wild type and D285K recordings were alternated.

**Supplemental Figure 1.**
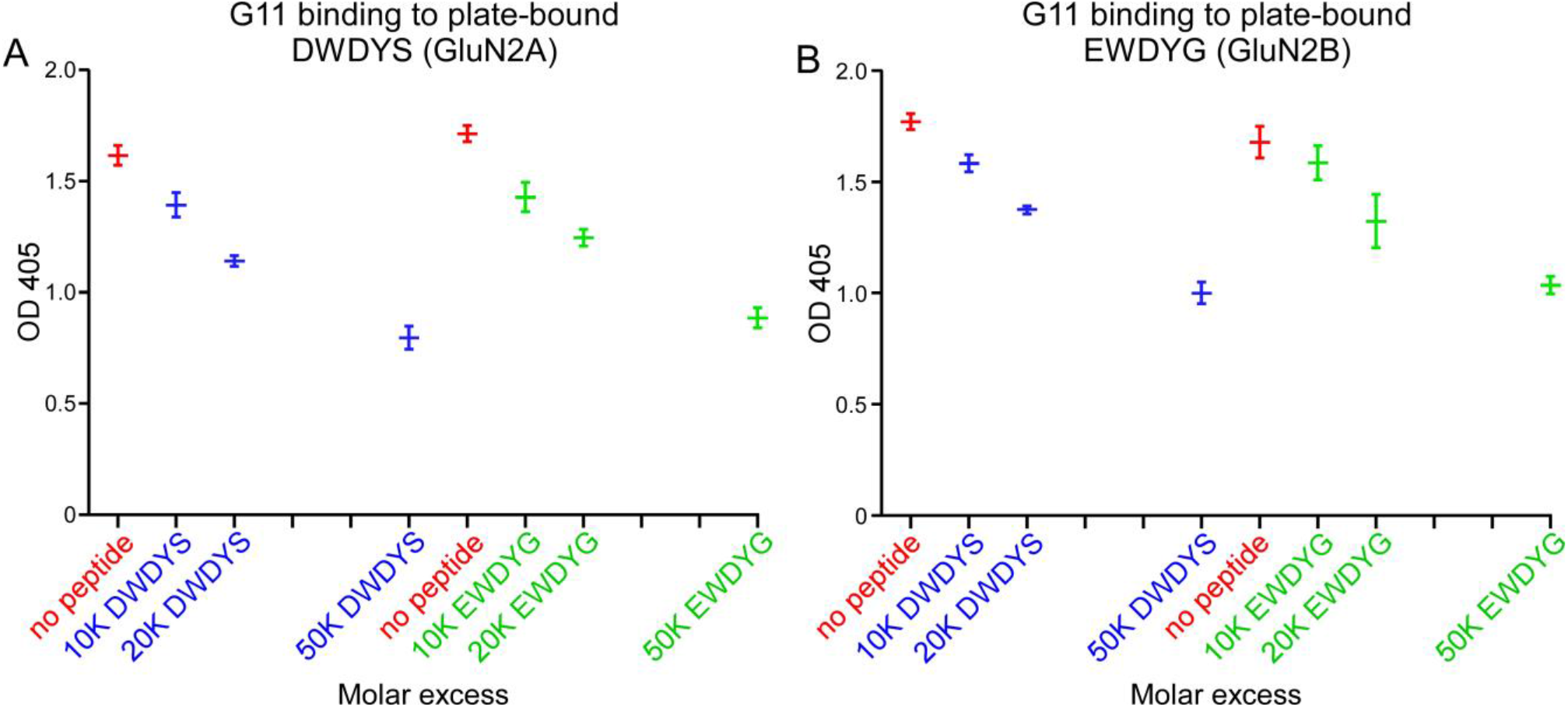
Competitive ELISAs for DNRAb binding to soluble GluN2A or GluN2B epitopes (relates to Figures 2B-2E). **(A & B)** Optical density (OD 405) values (mean ± SEM), measured by competitive ELISAs, for displacement of human DNRAb (G11) from plate-bound **(A)** DWDYS pentapeptide (GluN2A-specific) or **(B)** EWDYG (GluN2B-specific) by molar excess concentrations of the same soluble pentapeptides (n = 4 for each treatment condition).

**Supplemental Figure 2.**
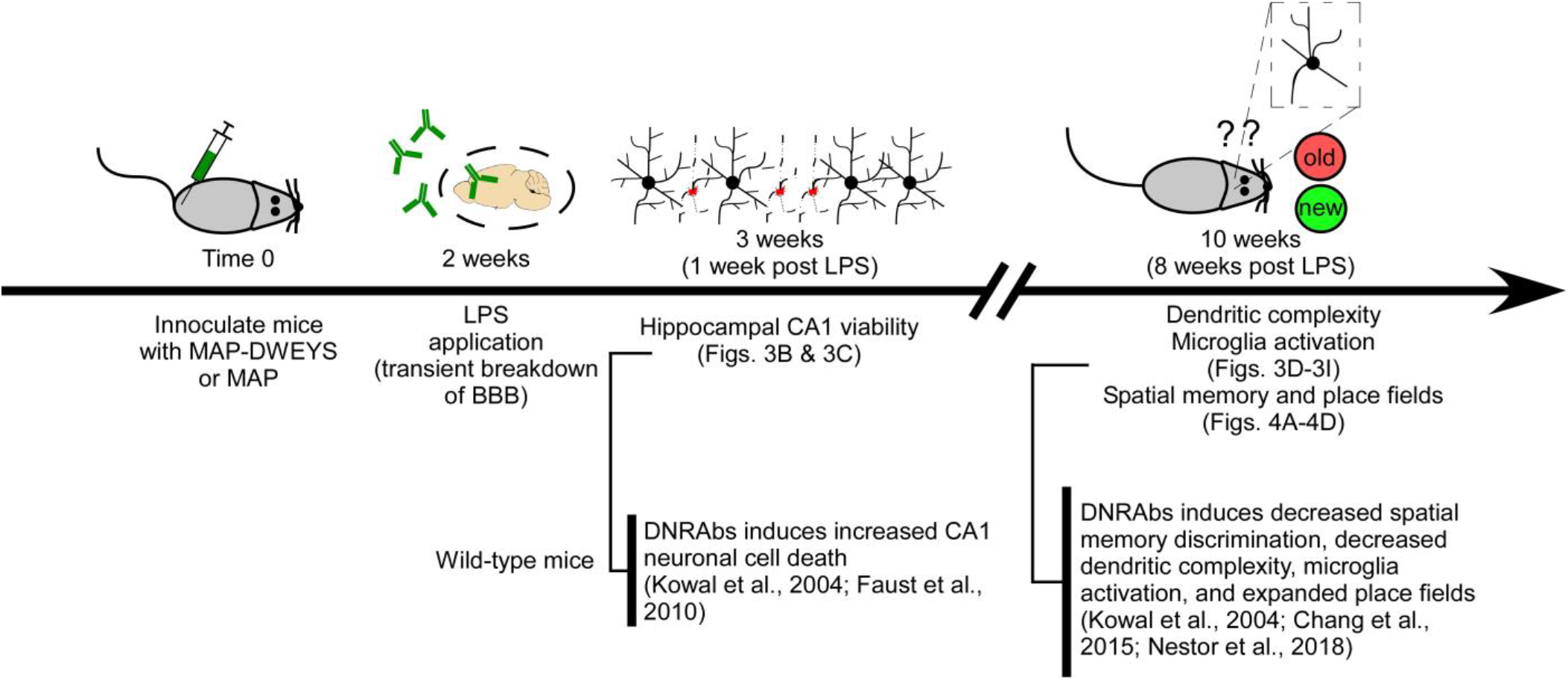
Diagram of the experimental protocol used for the NPSLE mouse model (relates to Figures 3 & 4). At time 0, mice were innoculated either with the DWEYS decapeptide multimerized on a polylysine backbone (MAP-DWEYS) or with the polylysine backbone alone (MAP). DWEYS is a mimetope of DNA and homologous to a sequence within the GluN2A and GluN2B amino-terminal domain. Immunization of wildtype mice with MAP-DWEYS induces production of DNRAbs (DNRAb+ mice) whereas MAP alone (control) does not^1^. Two weeks later, following two booster immunizations, mice were given lipopolysaccharide (LPS) to allow transient access of antibodies to the hippocampus^2, 3^. One week after the LPS treatment subsets of mice were sacrificed for histological characterization of CA1 pyramidal cell viability (Figs. 3B & 3C)^2, 4^. At this time point following LPS treatment, DNRAbs are still present in the hippocampus^2, 5^. Two weeks post-LPS, DNRAb levels are not detectable^5^. Eight weeks post LPS, subsets of animals were tested for hippocampal anatomy (CA1 dendritic complexity and microglia activation are altered in treated wild-type mice^3^ (Figs. 3D-3I). Other mice were tested for spatial memory test and implanted with tetrodes to determine place field size (Figs 4A-4D).

### Use of LPS to permealize BBB

LPS by itself produces a systemic inflammatory response and neuroinflammatory effects leading to neuronal death and microglia activation^6, 7, 8^. However, for all of our *in vivo* experiments, we only make comparisons between DNRAb+ (experimental) mice and DNARb- (control) mice, with both of these groups being treated with LPS. In addition, LPS permealization localizes DNRAbs more in the hippocampus and related structures than in other brain regions^2, 9^. The major region of action of DNRAbs in patients is the hippocampus and related structures^10, 11^.

## SUPPLEMENTAL METHODS

### Animals

All protocols were IACUC approved.

#### Feinstein Institute

Mice (females, C57BL/6 strain) were housed at the Center for Comparative Physiology at the Feinstein Institute for Medical Research. All protocols were IACUC approved. Mice with deletion of the GluN2A subunit (*Grin2a^−/−^*) were kindly provided by Dr. M. Mishina (University of Tokyo). Animals with conditional knockout of the GluN2B subunit (*Grin2b*^fl/fl^*;Camk2a-cre*) were bred in house by crossing homozygous floxed GluN2B mice (*Grin2b*^fl/fl^, kind gift from Prof. Dr. H. Monyer) with a Cre line driven by the CaMKIIα promoter (*Camk2a-cre* mice, B6.Cg-Tg(Camk2a-Cre)T29-1Stl/J, stock no: 005359, Jackson labs). Both groups were further bred on an H2d^+/+^ background in order to allow an antibody response to immunization. Immunization with MAP-core and MAP-DWEYS peptide was previously described ^1^. The first immunization was given in Complete Freund’s Adjuvant (Becton, Dickinson, and Company, 263810), with two boosters at 2 and 4 weeks in Incomplete Freund’s Adjuvant (Becton, Dickinson, and Company, 263910). At 2 weeks following the final immunization, mice received two intraperitoneal injections of LPS (6 mg/kg; Millipore-Sigma, L4524) 48 hours apart administered with a 500 μl intraperitoneal injection of sterile saline. Mice received water and food *ad libitum*.

#### Stony Brook

All animal procedures were approved by the institutional animal care and usage committee (IACUC) at Stony Brook University and were in concordance with the guidelines established by the National Institutes of Health. Mice (C57BL/6) were bred in house for primary cultures of hippocampal neurons at P0-P1 and glia/astrocyte feeder layers at P2-P4. Mice received water and food *ad libitum*.

### *In vitro* cell culture and transfection

Human embryonic kidney 293 (HEK293) cells were grown in Dulbecco’s modified Eagle’s medium (DMEM), supplemented with 10% FBS, for 24 h before transfection. Human NMDAR-encoding cDNA constructs (Table 1), were co-transfected into HEK293 cells along with a separate peGFP-Cl construct at a ratio of 4:4:1 (N1:N2:eGFP) for macroscopic recordings, and at a ratio for 4:1.5:1 for single channel recordings using X-tremeGene HP (Roche, 06-366). Triheteromeric NMDAR-expressing constructs (Table 1), all derived from rat, were a gift from K. Hansen and transfection with these constructs were performed as described previously ^2^. HEK293 cells were bathed in medium containing the GluN2 competitive antagonist DL-2-amino-5-phosphopentanoic acid (APV, 100 μM, Tocris), magnesium (100 μM), and the GluN1 competitive antagonist 6,7-dichlorokynuric acid (DCKA, 100 μM, Tocris). All experiments were performed 24-48 h post-transfection. Point mutations were introduced in the various GluN2A subunits using site-directed mutagenesis (SDM) as previously described ^3^

**Table 1.** Plasmid constructs used for heterologous expression of NMDARs in HEK293 cells.

| Construct | Species | Description |
| --- | --- | --- |
| pCI-neo-hGluN1a | Human | Wild-type receptor subunit |
| pCI-neo-hGluN2A | Human | Wild-type receptor subunit |
| pCI-neo-hGluN2B | Human | Wild-type receptor subunit |
| pCI-neo-2xATG-rGluN1a-GFP | Rat | Wild-type receptor subunit with bicistronic expression of eGFP (gift from K. Hansen) |
| pCI-neo-rGluN2A-C1-L4 | Rat | Tri-heteromeric modified subunit (Hansen et al., 2014) |
| pCI-neo-rGluN2A-C2-L4 | Rat | Tri-heteromeric modified subunit |
| pCI-neo-rGluN2B-AC1-L4 | Rat | Tri-heteromeric modified subunit |
| pCI-neo-rGluN2B-AC2-L4 | Rat | Tri-heteromeric modified subunit |

### *In vitro* macroscopic recordings

Macroscopic currents in the whole-cell mode were recorded at room temperature (20–23°C) using an EPC-10 amplifier with PatchMaster software (HEKA), digitized at 10 kHz and low-pass filtered at 2.9 kHz (−3 dB) using an 8 pole low pass Bessel filter ^4^. Patch microelectrodes were filled with an intracellular solution (mM): 140 KCl, 10 HEPES, 1 BAPTA, 4 Mg^2+^-ATP, 0.3 Na^+^-GTP, pH 7.3 (KOH), 297 mOsm (sucrose). Our standard extracellular solution consisted of (mM): 150 NaCl, 2.5 KCl, and 10 HEPES, pH 7.2 (NaOH). Pipettes had resistances of 2–6 MΩ when filled with the pipette solution and measured in the standard Na^+^ external solution. We did not use series resistance compensation nor did we correct for junction potentials. Currents were measured within 10 min of going whole cell.

External solutions were applied using a piezo-driven double barrel application system. Prior to use, we incubated the application system with 2% BSA (1xPBS) for two hours at 21–24°C to minimize non-specific antibody binding. For agonist application, one barrel contained the external solution +0.1 mM glycine, whereas the other barrel contained both 0.1 mM glycine and 1 mM glutamate. For display, NMDAR currents were digitally refiltered at 500 Hz and resampled at 1 kHz.

### *In vitro* single channel recordings

Single channel currents were recorded in the on-cell configuration at 20–23°C using an integrating patch clamp amplifier (Axopatch 200B, Molecular Devices), analog filtered at 10 kHz (four-pole Bessel filter), and digitized between 25 and 50 kHz (ITC-16 interfaced with PatchMaster, HEKA) ^5^.

Patch pipettes (thick-wall, borosilicate, Sutter Instruments) were pulled and fire-polished achieving resistances between 10 and 20 MΩ when measured in the bath. At −100 mV, seal resistance ranged between 2 and 20 GΩ. For cell-attached recordings, patch pipettes were filled with the standard bath solution as well as 1 mM glutamate and 0.1 mM glycine. 0.05 mM EDTA was added to minimize gating effects of divalents. Inward currents were elicited by applying a pipette potential of +100 mV. Recordings of hGluN1/hGluN2A or hGluN1/hGluN2B, either in the presence of absence of DNRAbs consisted of long clusters of activity separated by seconds-long periods of inactivity, simplifying detection of several channels in the patch. For these recordings the relatively high equilibrium open probability (eq. Po) and duration of recordings (≈10,000 to 180,000 per recording) indicated that we were recording from single-channel patches.

For 10 μg/ml antibody, our standard bath (pipette) solution consisted of 150 mM NaCl, 10 mM HEPES (pH 7.2, NaOH). For 100 μg/ml antibody, our standard bath had a decreased concentration of NaCl to 140 mM to compensate for the osmotic force generated by the antibodies in solution.

### Immunocytochemistry

HEK293T cells were used for immunocytochemistry (ICC). Transfections were performed in the same manner as described above, without GFP co-transfection. Cells were fixed with 4% paraformaldehyde 48 hours post-transfection, washed three times with 1xPBS, and then blocked with 2% bovine serum albumin (BSA) (Sigma, A9647) at room temperature. Incubation of primary antibody, either IgG control B1 (20 μg/mL) or human DNRAb, G11 (20 μg/mL) diluted in 2% BSA was done overnight at 4°C. Following three washes with 1xPBS, labeling was performed with secondary goat anti-human Alexa 488-conjugated antibodies (1:1000, ThermoFisher,A-11013) at room temperature. Coverslips were mounted with ProLong® Diamond Anti-fade mountant w/DAPI (ThermoFisher, P36962), and imaged with an Olympus FV-1000 confocal microscope. Five fields (212 × 212 μm, 60x oil) were taken per coverslip. Mean corrected fluorescence of fields were determined on ImageJ with spectral deconvolution plugin as previously described ^6, 7^ and ROI background correction.

For primary hippocampal neurons used for ICC, cultures were made according to methods as previously described ^8^ with minor modifications. Glia and astrocyte harvested from P2 mice ^9^ were seeded at 10-14 days prior to dissection. On day of dissection, glia/astrocyte cultures had media replaced with plating media, consisting of 10% (v/v) Fetal Bovine Serum (FBS) (ThermoFisher, 16140071), 0.45% (w/v) glucose, 1 mM sodium pyruvate, 2 mM glutamine, 100 U/mL penicillin/streptomycin in Basal Medium Eagle (ThermoFisher, 21010046). Hippocampi from P0-P1 C57/BL6 mice pips were dissected in ice-cold dissection media, consisting of 0.11 mg/mL sodium pyruvate, 0.1% glucose, 10 mM HEPES in HBSS, Ca^2+^ and Mg^2+^-free (ThermoFisher, 14175095) followed by 0.25% (v/v) trypsin-EDTA digestion for 20 min at 37°C. DNase I (Sigma, DN25) was added shortly after, and incubated for 5 min at RT. Following 2 washes with plating media, tissue was triturated with a FBS-coated pipette tip 10-12 times. Cells in suspension were counted with a hemocytometer, with approximately 150,000 cells added to 18 mm glass coverslips coated with poly-D-lysine in 12-well plates. Culture was incubated in plating medium for 2 hours at 37°C to allow for cells to adhere to coverslip. Coverslips were then flipped onto glia/astrocyte cultures, and cells were allowed to grow overnight in plating media, then switched with maintenance medium, consisting of 2% B-27 supplement (ThermoFisher, 17504044), 2 mM glutamine, 100 U/mL penicillin/streptomycin. On DIV3, 5 μM of cytosine arabinoside (Sigma, C1768) was added to cultures to inhibit mitosis of non-neuronal cells. Thereafter, half of maintenance media was replaced every 3 days until pharmacological treatment at DIV14.

Pharmacological treatment of primary hippocampal neurons occurred at DIV14. Final concentrations of antagonists were 3 μM: TCN-201 (AdooQ Bioscience, A11947), MPX-004 (Alomone Labs, M280), and Ifenprodil (Sigma, I2892). These were added along with the DNRAb (G11) at 10 μg/mL, in maintenance media to neurons. DMSO was used as vehicle control for both control (B1) and DNRAb (G11) conditions.

ICC was performed 24 hours post treatment on DIV15. Coverslips were washed with 1xPBS three times and then incubated in 4% paraformaldehyde(v/v)/4% sucrose (w/v) for 15 min. Coverslips were then washed again with 1xPBS three times and were blocked/permeabilized with antibody blocking solution, consisting of 2% (w/v) BSA, 0.25% (v/v) Triton-X100 in 1xPBS, for 60 min at RT. Coverslips were then incubated in primary antibodies, rabbit anti-activated caspase-3 (1:250, Cell Signaling, 9661), and mouse anti-β tubulin III (Tubb3) (1:400, Millipore-Sigma, MAB1637), both in antibody blocking solution overnight in 4°C. Coverslips were then washed with 1xPBS three times, and incubated in fluorescence-conjugated secondary antibodies, Goat anti-rabbitAlexa-647 (1:1000, ThermoFisher, A21245) and Goat anti-mouse Alexa-488(1:1000, ThermoFisher, A21121) for 1 hour at RT. Coverslips were then mounted and imaged as described above. Number of activated caspase-3 neurons were averaged by 5 fields (212 × 212 μm, 60x oil) per biological replicate. Quantification and image analysis were done on ImageJ by researcher blinded to treatment conditions.

### ELISA

Peptides were coated on COSTAR plates (Corning, CLS3690) at 20 μg/ml, 25 μl/well in 0.1M NaHCO_3_ pH 8.6, at 4°C, overnight. The plate was washed once with 1x PBS containing 0.05 % Tween 100 (PBS-T) and blocked with 1% BSA/PBS, 50 μl/well, for 1hr at 37°C. G11 antibody was used at 7.5 μg/ml which was on the linear range of binding curve established prior to the inhibition. Antibody was pre-incubated with a wide range of the inhibitory peptide in 1% BSA/PBS for 1 hr at RT and added to each well. The assay was performed twice with triplicates measurements. The well without the inhibitor peptide served as positive control. After incubation at 37°C for 1 hr, the plate was washed 6 times with PBS-T buffer. Goat anti-human secondary antibody labeled with alkaline phosphatase (IgG-AP) was added at 1:1000 dilution in 25 μl/well of 1% BSA/PBS followed by incubation at 37°C for 1 hr and by washing with PBS-T buffer. The ELISA was developed using AP substrate (Millipore-Sigma), in 50 μl/well of a buffer containing MgCl_2_ (1 mM final), Na_2_CO_3_ (25 mM final) and NaHCO_3_ (2.5 mM final). The reading was done at 405 μm using a multilabel counter from Perkin-Elmer.

### Immunohistochemistry

Mice were anesthetized with 100 μl of Euthasol (Virbac) prior to perfusion with 0.9% sodium chloride, 0.5% sodium nitrite, and 0.1% heparin, followed by 4% paraformaldehyde (PFA) in 0.1 M phosphate buffer (PB) as before^10^. Brains were extracted, fixed in 4% PFA for 2 hours and transferred to 30% sucrose, blocked and sliced (40 μm). Brains were blocked in a coronal stainless steel template and a 4 mm slab of tissue was cut from approximately Bregma −0.94 wewas sampled at a thickness of 40 microns over the next 1600 to 1900 microns.

For cresyl violet staining, sections were mounted, dehydrated, rehydrated and stained in cresyl violet for 3 min. Sections were dried, dehydrated, cleared (Histoclear II) and coverslipped (Permount, Fisher Scientific) prior to imaging on an AxiophotZ1 microscope (Zeiss). For immunohistochemistry, sections were washed (0.1 M phosphate buffered saline, PBS), permeabilized (0.2% Triton X-100 in 1% BSA in 0.1 M PBS), blocked (1% BSA in 0.1 M PBS for 60 minutes), and stained with primary antibody overnight at 4°C in 1% BSA. Primary antibodies were Iba1 (1:500, Wako Chemicals, 019-1 9741), CD68 (1:500, Bio-Rad, mca1957). On the following day, samples were washed (0.1 M PBS), incubated with secondary antibody (0.1 M PB for 45 min). Secondary antibodies included Donkey anti-rat Alexa Fluor 488 (1:400, Life Technologies, A21208), Chicken anti-rabbit Alexa Fluor 594 (1:300, Life Technologies, A21442). Sections underwent additional washes and incubation in DAPI (0.5 μg/mL in 0.1 M PB) and mounted with Cytoseal 60 (Thermo-Scientific) and cover-slipped.

Tissue was imaged on an AxioImager Z1 microscope or LSM 880 confocal microscope using super resolution Airy scan parameters (Zeiss). Microglia and colocalization with CD-68 were quantified using Zen2 Blue software as previously described^10, 11^. Microglia were quantitated per 106.4 μm^2^ box that was placed in 6 regions of interest across the stratum radiatum of CA1 using identical Airy scan acquisition parameters (40x oil, Airy = 2X, Z-stack = 10). Images of individual microglia were transferred to Neurolucida 360 (MBF Bioscience) for quantitation. We analyzed three tissue sections; the periodicity was one in four 40 μm sections across the dorsal CA1 hippocampus, there were three animals in each group, and 8-12 microglia per animal. All raw measurements were compiled for cumulative probability distributions and analyzed by Kolmogorov-Smirnov non-parametric statistics. The mean results for each Sholl dimension from the soma were plotted for each group and displayed in a standard fashion.

### Behavioral assessment and *in vivo* electrophysiology

Mice were handled for 15 min per day for 3 days before they were tested behaviorally. The object place memory (OPM) task was performed as described^10, 12^. The apparatus consisted of a square chamber (40 cm on the side, 40 cm high), with the walls painted grey. Animals were familiarized to the empty chamber (3 sessions of 15 min each). For OPM testing, mice underwent the following sequence: empty chamber (10 min), home cage (10 min), sample phase in which the chamber had two objects located at the center of the NW and NE sectors (5 min), home cage (10 min), choice phase in which the chamber had the same objects but one of them was moved from NE to the center of the SE sector (5 min). The discrimination ratio was calculated during the choice phase by dividing time spent exploring the moved object minus the time spent exploring the static object by the time exploring the objects combined^12^. Data were collected and analyzed using EthoVision XT (Noldus Information Technologies).

The analysis of place cells in the dorsal CA1 region of the hippocampus was completed as previously described^12, 13^. A mouse was anesthetized with 0.25% isoflorane and implanted with a 16 channel multi-electrode array containing 4 tetrodes. After recovery, single unit firing was recorded using Cheetah software (Neuralynx) while the animal explored a square chamber (40 cm on the side) in a schedule of four exploration runs (15 min) separated by three rest sessions (5 min) in the homecage. Recordings were repeated over 2 consecutive days. Acquired data were analyzed using Spike2 (version 8, Cambridge Electronic Design), NeuroExplorer (version 5, Nex Technologies), and Matlab.

## REFERENCES

1. Tsokos GC. Systemic lupus erythematosus. N Engl J Med 365, 2110–2121 (2011).

2. Chang EH, et al. Selective Impairment of Spatial Cognition Caused by Autoantibodies to the N-Methyl-D-Aspartate Receptor. EBioMedicine 2, 755–764 (2015).

3. Kowal C, et al. Human lupus autoantibodies against NMDA receptors mediate cognitive impairment. Proc Natl Acad Sci U S A 103, 19854–19859 (2006).

4. Mackay M, etal. Metabolic and microstructural alterations in the SLE brain correlate with cognitive impairment. JCI Insight 4, (2019).

5. Hanly JG, et al. Prospective analysis of neuropsychiatric events in an international disease inception cohort of patients with systemic lupus erythematosus. Ann Rheum Dis 69, 529–535 (2010).

6. Unterman A, Nolte JE, Boaz M, Abady M, Shoenfeld Y, Zandman-Goddard G. Neuropsychiatric syndromes in systemic lupus erythematosus: a meta-analysis. Semin Arthritis Rheum 41, 1–11 (2011).

7. Borowoy AM, et al. Neuropsychiatric lupus: the prevalence and autoantibody associations depend on the definition: results from the 1000 faces of lupus cohort. Semin Arthritis Rheum 42, 179–185 (2012).

8. DeGiorgio LA, Konstantinov KN, Lee SC, Hardin JA, Volpe BT, Diamond B. A subset of lupus anti-DNA antibodies cross-reacts with the NR2 glutamate receptor in systemic lupus erythematosus. Nat Med 7, 1189–1193 (2001).

9. Arinuma Y, Yanagida T, Hirohata S. Association of cerebrospinal fluid anti-NR2 glutamate receptor antibodies with diffuse neuropsychiatric systemic lupus erythematosus. Arthritis Rheum 58, 1130–1135 (2008).

10. Faust TW, et al. Neurotoxic lupus autoantibodies alter brain function through two distinct mechanisms. Proc Natl Acad Sci U S A 107, 18569–18574 (2010).

11. Kowal C, et al. Cognition and immunity; antibody impairs memory. Immunity 21, 179–188 (2004).

12. Kapadia M, et al. Effects of sustained i.c.v. infusion of lupus CSF and autoantibodies on behavioral phenotype and neuronal calcium signaling. Acta Neuropathol Commun 5, 70 (2017).

13. Gono T, et al. Anti-NR2A antibody as a predictor for neuropsychiatric systemic lupus erythematosus. Rheumatology (Oxford) 50, 1578–1585 (2011).

14. Tay SH, Fairhurst AM, Mak A. Clinical utility of circulating anti-N-methyl-d-aspartate receptor subunits NR2A/B antibody for the diagnosis of neuropsychiatric syndromes in systemic lupus erythematosus and Sjogren’s syndrome: An updated meta-analysis. Autoimmun Rev 16, 114–122 (2017).

15. Zhang J, et al. Identification of DNA-reactive B cells in patients with systemic lupus erythematosus. J Immunol Methods 338, 79–84 (2008).

16. Zhang J, Jacobi AM, Wang T, Berlin R, Volpe BT, Diamond B. Polyreactive autoantibodies in systemic lupus erythematosus have pathogenic potential. J Autoimmun 33, 270–274 (2009).

17. Huerta PT, Kowal C, DeGiorgio LA, Volpe BT, Diamond B. Immunity and behavior: antibodies alter emotion. Proc Natl Acad Sci U S A 103, 678–683 (2006).

18. Appenzeller S, Carnevalle AD, Li LM, Costallat LT, Cendes F. Hippocampal atrophy in systemic lupus erythematosus. Ann Rheum Dis 65, 1585–1589 (2006).

19. Nestor J, et al. Lupus antibodies induce behavioral changes mediated by microglia and blocked by ACE inhibitors. J Exp Med, (2018).

20. Traynelis SF, et al. Glutamate receptor ion channels: structure, regulation, and function. Pharmacol Rev 62, 405–496 (2010).

21. Paoletti P, Bellone C, Zhou Q. NMDA receptor subunit diversity: impact on receptor properties, synaptic plasticity and disease. Nat Rev Neurosci 14, 383–400 (2013).

22. Hansen KB, et al. Structure, function, and allosteric modulation of NMDA receptors. J Gen Physiol 150, 1081–1105 (2018).

23. Wang L, et al. Female mouse fetal loss mediated by maternal autoantibody. J Exp Med 209, 1083–1089 (2012).

24. Talukder I, Wollmuth LP. Local constraints in either the GluN1 or GluN2 subunit equally impair NMDA receptor pore opening. J Gen Physiol 138, 179–194 (2011).

25. Amin JB, Leng X, Gochman A, Zhou HX, Wollmuth LP. A conserved glycine harboring disease-associated mutations permits NMDA receptor slow deactivation and high Ca(2+) permeability. Nat Commun 9, 3748 (2018).

26. Rauner C, Kohr G. Triheteromeric NR1/NR2A/NR2B receptors constitute the major N-methyl-D-aspartate receptor population in adult hippocampal synapses. J Biol Chem 286, 7558–7566 (2011).

27. Tovar KR, McGinley MJ, Westbrook GL. Triheteromeric NMDA receptors at hippocampal synapses. J Neurosci 33, 9150–9160 (2013).

28. Hansen KB, Ogden KK, Yuan H, Traynelis SF. Distinct functional and pharmacological properties of Triheteromeric GluN1/GluN2A/GluN2B NMDA receptors. Neuron 81, 1084–1096 (2014).

29. Stroebel D, Carvalho S, Paoletti P. Functional evidence for a twisted conformation of the NMDA receptor GluN2A subunit N-terminal domain. Neuropharmacology 60, 151–158 (2011).

30. Romero-Hernandez A, Simorowski N, Karakas E, Furukawa H. Molecular Basis for Subtype Specificity and High-Affinity Zinc Inhibition in the GluN1-GluN2A NMDA Receptor Amino-Terminal Domain. Neuron 92, 1324–1336 (2016).

31. Volkmann RA, et al. MPX-004 and MPX-007: New Pharmacological Tools to Study the Physiology of NMDA Receptors Containing the GluN2A Subunit. PLoS One 11, e0148129 (2016).

32. Gono T, et al. NR2-reactive antibody decreases cell viability through augmentation of Ca(2+) influx in systemic lupus erythematosus. Arthritis Rheum 63, 3952–3959 (2011).

33. Sakimura K, et al. Reduced hippocampal LTP and spatial learning in mice lacking NMDA receptor epsilon 1 subunit. Nature 373, 151–155 (1995).

34. Kutsuwada T, et al. Impairment of suckling response, trigeminal neuronal pattern formation, and hippocampal LTD in NMDA receptor epsilon 2 subunit mutant mice. Neuron 16, 333–344 (1996).

35. von Engelhardt J, et al. Contribution of hippocampal and extra-hippocampal NR2B-containing NMDA receptors to performance on spatial learning tasks. Neuron 60, 846–860 (2008).

36. Putterman C, Diamond B. Immunization with a peptide surrogate for double-stranded DNA (dsDNA) induces autoantibody production and renal immunoglobulin deposition. J Exp Med 188, 29–38 (1998).

37. Larkin AE, et al. Blockade of NMDA receptors pre-training, but not post-training, impairs object displacement learning in the rat. Brain Res 1199, 126–132 (2008).

38. Faust TW, Robbiati S, Huerta TS, Huerta PT. Dynamic NMDAR-mediated properties of place cells during the object place memory task. Front Behav Neurosci 7, 202 (2013).

39. Ekstrom AD, Meltzer J, McNaughton BL, Barnes CA. NMDA receptor antagonism blocks experience-dependent expansion of hippocampal “place fields”. Neuron 31, 631–638 (2001).

40. McHugh TJ, Blum KI, Tsien JZ, Tonegawa S, Wilson MA. Impaired hippocampal representation of space in CA1-specific NMDAR1 knockout mice. Cell 87, 1339–1349 (1996).

41. Cacucci F, Yi M, Wills TJ, Chapman P, O’Keefe J. Place cell firing correlates with memory deficits and amyloid plaque burden in Tg2576 Alzheimer mouse model. Proc Natl Acad Sci U S A 105, 7863–7868 (2008).

42. Bertsias GK, Boumpas DT. Pathogenesis, diagnosis and management of neuropsychiatric SLE manifestations. Nat Rev Rheumatol 6, 358–367 (2010).

43. Fragoso-Loyo H, et al. Serum and cerebrospinal fluid autoantibodies in patients with neuropsychiatric lupus erythematosus. Implications for diagnosis and pathogenesis. PLoS One 3, e3347 (2008).

44. Omdal R, Brokstad K, Waterloo K, Koldingsnes W, Jonsson R, Mellgren SI. Neuropsychiatric disturbances in SLE are associated with antibodies against NMDA receptors. Eur J Neurol 12, 392–398 (2005).

45. Sun W, Hansen KB, Jahr CE. Allosteric Interactions between NMDA Receptor Subunits Shape the Developmental Shift in Channel Properties. Neuron 94, 58–64 e53 (2017).

46. Lu W, Du J, Goehring A, Gouaux E. Cryo-EM structures of the triheteromeric NMDA receptor and its allosteric modulation. Science 355, (2017).

47. Hackos DH, et al. Positive Allosteric Modulators of GluN2A-Containing NMDARs with Distinct Modes of Action and Impacts on Circuit Function. Neuron 89, 983–999 (2016).

48. Yi F, et al. Structural Basis for Negative Allosteric Modulation of GluN2A-Containing NMDA Receptors. Neuron 91, 1316–1329 (2016).

49. Tajima N, et al. Activation of NMDA receptors and the mechanism of inhibition by ifenprodil. Nature 534, 63–68 (2016).

50. Zhu S, et al. Mechanism of NMDA Receptor Inhibition and Activation. Cell 165, 704–714 (2016).

51. Esmenjaud JB, et al. An inter-dimer allosteric switch controls NMDA receptor activity. EMBO J, (2018).

52. Gielen M, Siegler Retchless B, Mony L, Johnson JW, Paoletti P. Mechanism of differential control of NMDA receptor activity by NR2 subunits. Nature 459, 703–707 (2009).

53. Ogden KK, Traynelis SF. New advances in NMDA receptor pharmacology. Trends Pharmacol Sci 32, 726–733 (2011).

54. Chen HS, Lipton SA. The chemical biology of clinically tolerated NMDA receptor antagonists. J Neurochem 97, 1611–1626 (2006).

55. Yelshansky MV, Sobolevsky AI, Jatzke C, Wollmuth LP. Block of AMPA receptor desensitization by a point mutation outside the ligand-binding domain. J Neurosci 24, 4728–4736 (2004).

56. Alsaloum M, Kazi R, Gan Q, Amin J, Wollmuth LP. A Molecular Determinant of Subtype-Specific Desensitization in Ionotropic Glutamate Receptors. J Neurosci 36, 2617–2622 (2016).

57. Kazi R, Dai J, Sweeney C, Zhou HX, Wollmuth LP. Mechanical coupling maintains the fidelity of NMDA receptor-mediated currents. Nat Neurosci 17, 914–922 (2014).

58. Beaudoin GM, 3rd, et al. Culturing pyramidal neurons from the early postnatal mouse hippocampus and cortex. Nat Protoc 7, 1741–1754 (2012).

## References used in Supplemental Material

1. Putterman C, Diamond B. Immunization with a peptide surrogate for double-stranded DNA (dsDNA) induces autoantibody production and renal immunoglobulin deposition. J Exp Med 188, 29–38 (1998).

2. Kowal C, et al. Cognition and immunity; antibody impairs memory. Immunity 21, 179–188 (2004).

3. Nestor J, et al. Lupus antibodies induce behavioral changes mediated by microglia and blocked by ACE inhibitors. J Exp Med, (2018).

4. Faust TW, et al. Neurotoxic lupus autoantibodies alter brain function through two distinct mechanisms. Proc Natl Acad Sci U S A 107, 18569–18574 (2010).

5. Chang EH, et al. Selective Impairment of Spatial Cognition Caused by Autoantibodies to the N-Methyl-D-Aspartate Receptor. EBioMedicine 2, 755–764 (2015).

6. Sternberg EM. Neural-immune interactions in health and disease. J Clin Invest 100, 2641–2647 (1997).

7. Nolan Y, Vereker E, Lynch AM, Lynch MA. Evidence that lipopolysaccharide-induced cell death is mediated by accumulation of reactive oxygen species and activation of p38 in rat cortex and hippocampus. Exp Neurol 184, 794–804 (2003).

8. Vereker E, Campbell V, Roche E, McEntee E, Lynch MA. Lipopolysaccharide inhibits long term potentiation in the rat dentate gyrus by activating caspase-1. J Biol Chem 275, 26252–26258 (2000).

9. Huerta PT, Kowal C, DeGiorgio LA, Volpe BT, Diamond B. Immunity and behavior: antibodies alter emotion. Proc Natl Acad Sci U S A 103, 678–683 (2006).

10. Appenzeller S, Carnevalle AD, Li LM, Costallat LT, Cendes F. Hippocampal atrophy in systemic lupus erythematosus. Ann Rheum Dis 65, 1585–1589 (2006).

11. Mackay M, et al. Metabolic and microstructural alterations in the SLE brain correlate with cognitive impairment. JCI Insight 4, (2019).

## SUPPLEMENTAL METHODS REFERENCES

1. Kowal C, et al. Cognition and immunity; antibody impairs memory. Immunity 21, 179–188 (2004).

2. Hansen KB, Ogden KK, Yuan H, Traynelis SF. Distinct functional and pharmacological properties of Triheteromeric GluN1/GluN2A/GluN2B NMDA receptors. Neuron 81, 1084–1096 (2014).

3. Kazi R, Dai J, Sweeney C, Zhou HX, Wollmuth LP. Mechanical coupling maintains the fidelity of NMDA receptor-mediated currents. Nat Neurosci 17, 914–922 (2014).

4. Yelshansky MV, Sobolevsky AI, Jatzke C, Wollmuth LP. Block of AMPA receptor desensitization by a point mutation outside the ligand-binding domain. J Neurosci 24, 4728–4736 (2004).

5. Talukder I, Wollmuth LP. Local constraints in either the GluN1 or GluN2 subunit equally impair NMDA receptor pore opening. J Gen Physiol 138, 179–194 (2011).

6. Gammon ST, Leevy WM, Gross S, Gokel GW, Piwnica-Worms D. Spectral unmixing of multicolored bioluminescence emitted from heterogeneous biological sources. Anal Chem 78, 1520–1527 (2006).

7. Zurek-Biesiada D, Kedracka-Krok S, Dobrucki JW. UV-activated conversion of Hoechst 33258, DAPI, and Vybrant DyeCycle fluorescent dyes into blue-excited, green-emitting protonated forms. Cytometry A 83, 441–451 (2013).

8. Beaudoin GM, 3rd, et al. Culturing pyramidal neurons from the early postnatal mouse hippocampus and cortex. Nat Protoc 7, 1741–1754 (2012).

9. Kaech S, Banker G. Culturing hippocampal neurons. Nat Protoc 1, 2406–2415 (2006).

10. Nestor J, et al. Lupus antibodies induce behavioral changes mediated by microglia and blocked by ACE inhibitors. J Exp Med, (2018).

11. Schafer DP, et al. Microglia sculpt postnatal neural circuits in an activity and complement-dependent manner. Neuron 74, 691–705 (2012).

12. Chang EH, et al. Selective Impairment of Spatial Cognition Caused by Autoantibodies to the N-Methyl-D-Aspartate Receptor. EBioMedicine 2, 755–764 (2015).

13. Faust TW, et al. Neurotoxic lupus autoantibodies alter brain function through two distinct mechanisms. Proc Natl Acad Sci U S A 107, 18569–18574 (2010).

